# Life-cycle-related gene expression patterns in the brown algae

**DOI:** 10.1101/2025.04.25.649966

**Authors:** Pélagie Ratchinski, Olivier Godfroy, Benjamin Noel, Jean-Marc Aury, J. Mark Cock

## Abstract

Brown algae are important primary constituents of marine coastal ecosystems, characterised by complex life cycles and various levels of complex multicellular development. However, the molecular processes that underlie development and life cycle progression in the brown algae remain poorly understood. In this study, pairwise comparisons of gametophyte and sporophyte transcriptomes across ten diverse brown algal species showed that the total number of genes exhibiting generation-biased or generation-specific expression in each species was correlated with the degree of dimorphism between life cycle generations. However, analysis of gene ontology terms assigned to the generation-biased/generation-specific genes indicated that each generation (i.e. the sporophyte and the gametophyte) also has characteristic broad life-cycle-related features that have been conserved during evolution. A more detailed analysis of *Ectocarpus* species 7, identified progressive transcriptome changes over its entire life cycle with a particularly marked change in transcriptome composition during the first day of sporophyte development, characterised by downregulation of flagellar and transcription factor genes and upregulation of a subset of translation genes. Comparison with a similar transcriptomic time series for the evolutionarily-distant (about 250 My) brown alga *Dictyota dichotoma* indicated considerable conservation of co-expressed gene modules between the two species, particularly for modules that were enriched in genes assigned to evolutionarily-conserved functional categories. This study therefore identified broad life-cycle- and development-related patterns of gene expression that are conserved across the brown algae.

## Introduction

Brown algae are major primary components of diverse and widespread coastal ecosystems, often forming extensive underwater forests (Bringloe *et al*., 2020; Eger *et al*., 2023). The group includes large (up to 50 metres) multicellular organisms that rival many land plant species in complexity. From a developmental point of view, brown algae are particularly interesting because they evolved complex multicellularity independently of land plants and animals (Cock *et al*., 2010). Taxonomically, these seaweeds are grouped into the class Phaeophyceae, which is a major taxon within the stramenopiles, and therefore very distant, phylogenetically, from the Archaeplastida and the Opisthokonta lineages that gave rise to land plants and animals, respectively. Consequently, brown algae represent interesting alternative systems to investigate the evolution and molecular bases of developmental processes.

Most brown algae have haploid-diploid life cycles involving an alternation between two independent multicellular generations, the sporophyte and the gametophyte (Cock *et al*., 2014). These seaweeds are therefore capable of deploying two different multicellular developmental programs, each at the appropriate stage of their complex life cycles. The ecological functions of brown algal haploid-diploid life cycles can be difficult to study, particularly when one of the generations is microscopic. However, there is evidence, in kelps for example, that microscopic stages (gametophytes, but possibly also very young sporophyte stages) may have a survival function, allowing a population to persist when the macroscopic sporophyte stage is not present (Carney and Edwards, 2006). A study of populations of filamentous brown algae of the genus *Ectocarpus* in the field indicated that gametophytes grew epiphytically on the brown alga *Scytosiphon promiscuus* (then classified taxonomically as *Scytosiphon lomentaria*) in the spring, whereas sporophytes grew throughout the year on abiotic substrata. These observations indicate that the sporophyte may be the most important generation for persistence of the population in this genus (Couceiro *et al*., 2015). Note, however, that a principally asexual population of *Ectocarpus siliculosus* located at a second site inhabited both epiphytic and epilithic niches indicating that the relationship between life cycle generation and niche may be complex (Couceiro *et al*., 2015). The phenology of the brown alga *D. dichotoma* also appears to be complex with, for example, overlapping generations and sporophytes and gametophytes reported to occur simultaneously (Tronholm *et al*., 2008).

The relative sizes of the two life cycle generations varies considerably across brown algae, with either the sporophyte or the gametophyte being the largest generation (dimorphism) or the two generations having very similar sizes and morphologies (isomorphy). Transitions between life cycles with different degrees of dimorphism have occurred frequently during the evolution of the brown algae (Cock et al., 2014). Variations on this central sexual life cycle have been observed in culture for several species and these variations can provide insights into the mechanisms that control life cycle progression. For example, if *Ectocarpus* gametes fail to fuse with a gamete of the opposite sex to form a zygote, they are often capable of germinating parthenogenetically to produce a haploid partheno-sporophyte that is morphologically and functionally indistinguishable from a diploid sporophyte (Müller, 1967). The existence of these haploid sporophytes indicates that life cycle generation identity is not determined by ploidy in this species and is therefore presumably under genetic control.

The occurrence of two multicellular generations during the life cycles of most brown algae allows genetic screens to be carried out for mutations that cause switching between the development programs of the two generations, i.e. mutations that lead to deployment of the incorrect program at a specific point in the life cycle (Coelho and Cock, 2020). Using this approach, two three-amino-acid-loop-extension (TALE) homeodomain transcription factor (HD TF) genes, *OUROBOROS (ORO)* or *SAMSARA (SAM)*, have been show to be necessary for deployment of the sporophyte generation developmental program in *Ectocarpus* (Coelho *et al*., 2011; Arun *et al*., 2019). Mutations in either of these genes suppresses the ability of the alga to produce a functional sporophyte, leading to deployment of the gametophyte program at life cycle stages where a sporophyte should normally be produced. In addition, *Ectocarpus* sporophytes have been shown to produce a diffusible factor that can induce sporophyte development in gametophyte initial cells (Arun *et al*., 2013; Yao *et al*., 2021). The diffusible factor does not induce life cycle generation switching in *oro* or *sam* mutant gametophyte initial cells, indicating that it acts upstream of ORO and SAM (Arun *et al*., 2013, 2019).

The genetic pathways that underlie the deployment of the sporophyte and gametophyte developmental programs in brown algae are still poorly understood. Mutations in a small number of genes have been shown to lead to developmental defects in *Ectocarpus* (Macaisne *et al*., 2017; Godfroy *et al*., 2017, 2023) but a mutation in the *IMMEDIATE UPRIGHT* gene is the only lesion that specifically affects the development of one of the two generations (Peters *et al*., 2008). Transcriptomics has been used to identify genes that are differentially expressed between the two generations in *Ectocarpus* (Arun *et al*., 2019; Lipinska *et al*., 2019; Bourdareau *et al*., 2021). These transcriptomic analyses focused on comparisons of adult sporophyte and gametophyte thalli and have allowed the identification of several hundred genes that are differentially expressed between these two stages of development. However, the analyses were not designed to detect genes that are important during early development and do not provide any information about the kinetics of gene expression over development. In recent years, several detailed studies have generated gene expression data for multiple developmental stages or organs for several different brown algal species (Pearson *et al*., 2019; Shao *et al*., 2019; Shan *et al*., 2020; Zhang *et al*., 2021; Graf *et al*., 2022; Liang *et al*., 2023; Uji *et al*., 2024), including a transcriptomic analyses of early stages of *Dictyota dichotoma* sporophyte development (Bogaert *et al*., 2017). This latter study detected changes in the transcriptome less than one hour after fertilisation, providing evidence for *de novo* transcription in the zygote, and identified a set of genes with diverse functions that were differentially regulated during early sporophyte development.

In the current study, transcriptomic data, recently generated by the Phaeoexplorer project (Denoeud *et al*., 2024), was used to analyse differential gene expression between the adult sporophyte and gametophyte generations of a broad range of brown algae. The proportion of genes that showed generation-biased or generation-specific expression across these two stages was correlated with the level of dimorphism between the two generations indicating that the differentially expressed genes are primarily linked to morphological differentiation. However, analysis of gene ontology terms also identified underlying, life-cycle-related characteristics that have been conserved during evolution, with general classes of gene function being conserved across sporophytes of the ten species, and different general classes of gene function being conserved across the gametophytes. In addition, analysis of transcriptomic data corresponding to multiple stages of the *Ectocarpus* life cycle, generated by combining published RNA sequencing (RNA-seq) data with newly-generated timepoints, identified co-expression modules containing genes with specific expression patterns during life cycle progression. This analysis identified several modules that exhibited marked expression pattern changes during early sporophyte evolution, providing insights into key genetic events during this period. Comparison with a set of gene co-expression modules generated for *D. dichotoma* identified conserved features shared by these two distantly related brown algal species.

## Results

### Across the brown algal lineage, the proportion of generation-biased genes is correlated with the degree of life cycle dimorphism

RNA-seq data for the adult sporophyte and gametophyte generations of ten different brown algal species (Figure 1A, Table S1) were analysed to obtain a broad overview of generation-biased gene expression across the brown algae. The species analysed had markedly different haploid-diploid life cycles, ranging from sporophyte dominance, through sporophyte-gametophyte isomorphy to gametophyte dominance (Figure S1). As previously observed in a smaller-scale analysis that looked at four different brown algal species (Lipinska *et al*., 2019), the proportion of generation-biased/generation-specific genes (see Materials and Methods for definitions of generation-biased and generation-specific genes) identified in each species was positively correlated with the degree of intergenerational dimorphism (Figure 1B, S1, Table S2). Surprisingly, however, species with strongly dimorphic life cycles had greater numbers of both sporophyte- and gametophyte-biased/specific genes, indicating either that gene downregulation plays an important role in the establishment of dimorphic generations or that there is also cryptic complexification of the more morphologically simple generation.

**Figure 1.**
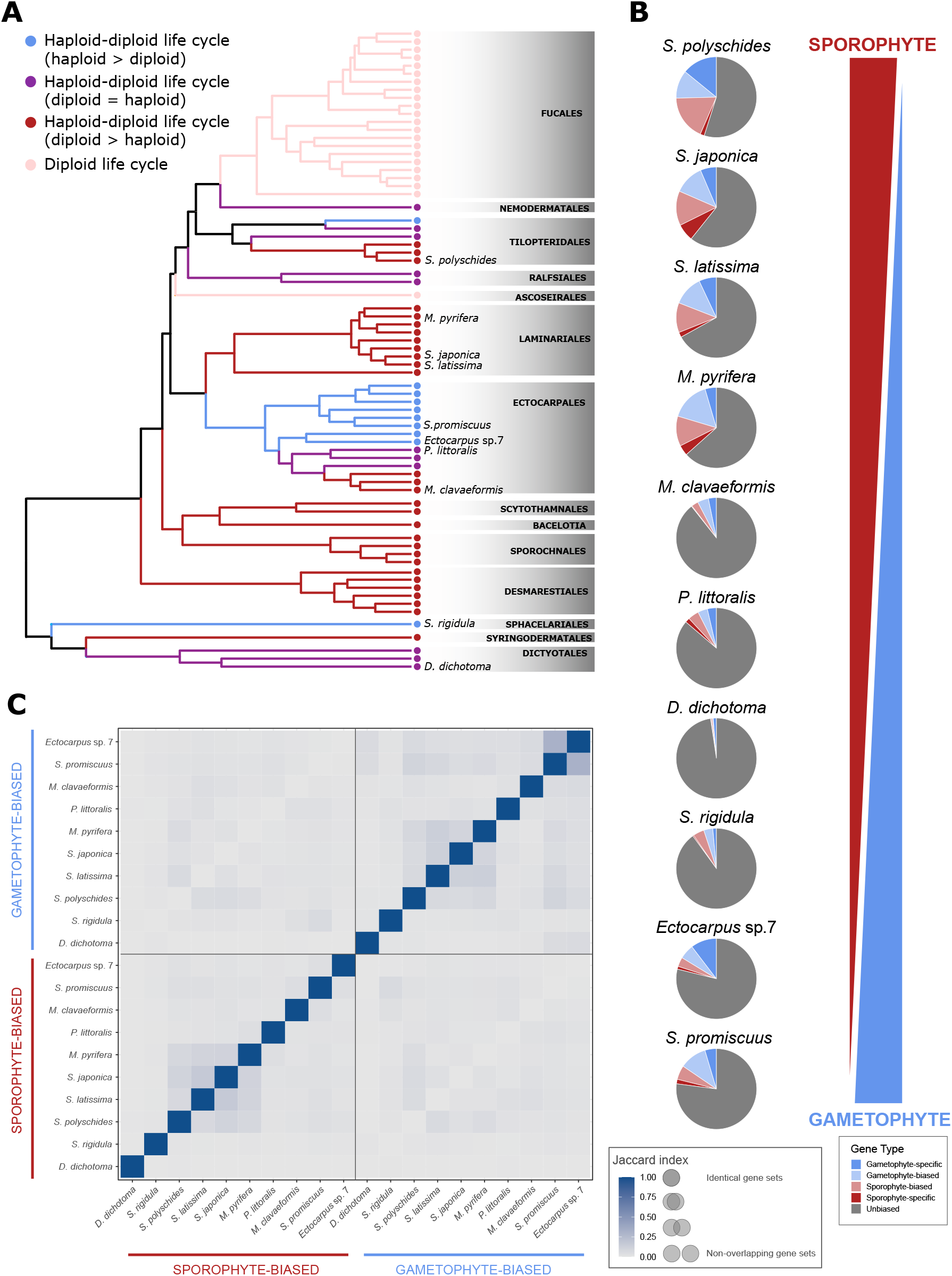
Generation-biased gene expression across the brown algae. **A**. Cladogram of the brown algae showing the ten species for which generation-biased gene expression was compared. Adapted from (Cock *et al*., 2014). **B**. Generation-biased gene expression in relation to life cycle dimorphism. Pie diagrams indicating the proportions of sporophyte-specific (dark red), sporophyte-biased (light red), gametophyte-specific (dark blue), gametophyte-biased (light blue) and unbiased (grey) genes in species with different haploid-diploid life cycles ranging from sporophyte-dominant through isomorphic to gametophyte-dominant (indicated by the red and blue wedges). sp., species. C. Overlaps (Jaccard index) between sets of gametophyte-biased/specific and sporophyte-biased/specific genes across the ten species.

We also noted that, depending on the species, only between 10% and 24% (Table S2) of genes were expressed during one generation but not the other (defined for this analysis as mean TPM ⌇1 for one generation but mean TPM <1 in the other generation), indicating that most genes are expressed during both generations of the life cycle. There is therefore considerable scope for many genes to have functions during both generations of the life cycle. In terms of selection, this observation indicates that most genes are potentially directly exposed to purifying selection during the haploid phase (where there is no effect of masking by a second allele of each locus) and few genes are expressed solely in a diploid context where masking would be most effective.

### The sporophyte and gametophyte generations express characteristic general functional categories of genes

To evaluate the degree of conservation of generation-biased/specific gene sets across the ten brown algal species an orthogroup analysis was carried out (Table S3) using Orthofinder (Emms and Kelly, 2019). Pairwise comparisons of generation-biased/specific genes that had orthologues in all ten species indicated that the sets of generation-biased/specific genes were poorly conserved across species, even for closely-related species such as the two *Saccharina* spp. (Figure 1C). None of the orthogroups exhibited conserved generation bias across all the ten species. The lack of conservation of generation-biased/specific gene sets across species is in line with observations made in the earlier study that compared four brown algal species (Lipinska *et al*., 2019). However, despite the poor conservation of generation-biased/specific expression at the individual gene level, analysis of enriched gene ontology terms across the entire generation-biased/specific gene sets from the ten studied brown algae identified distinct sets of enriched terms for both the sporophyte-biased/specific and the gametophyte-biased/specific genes (Figure 2) and the general classes of enriched gene ontology term were conserved for each individual generation across the ten species. For example, the sporophyte-biased/specific gene sets tended to be enriched in genes with predicted functions related to membrane transport and metabolism whereas the gametophyte-biased/specific gene sets tended to be enriched in genes with predicted functions related to flagella biosynthesis and function, protein biosynthesis and DNA-related functions (Figure 2). These analyses identified an underlying similarity between sporophyte generations across species and between gametophyte generations across species, and indicated that these generation-characteristic patterns of gene expression are present even in species where the two generations are isomorphic or nearly isomorphic.

**Figure 2.**
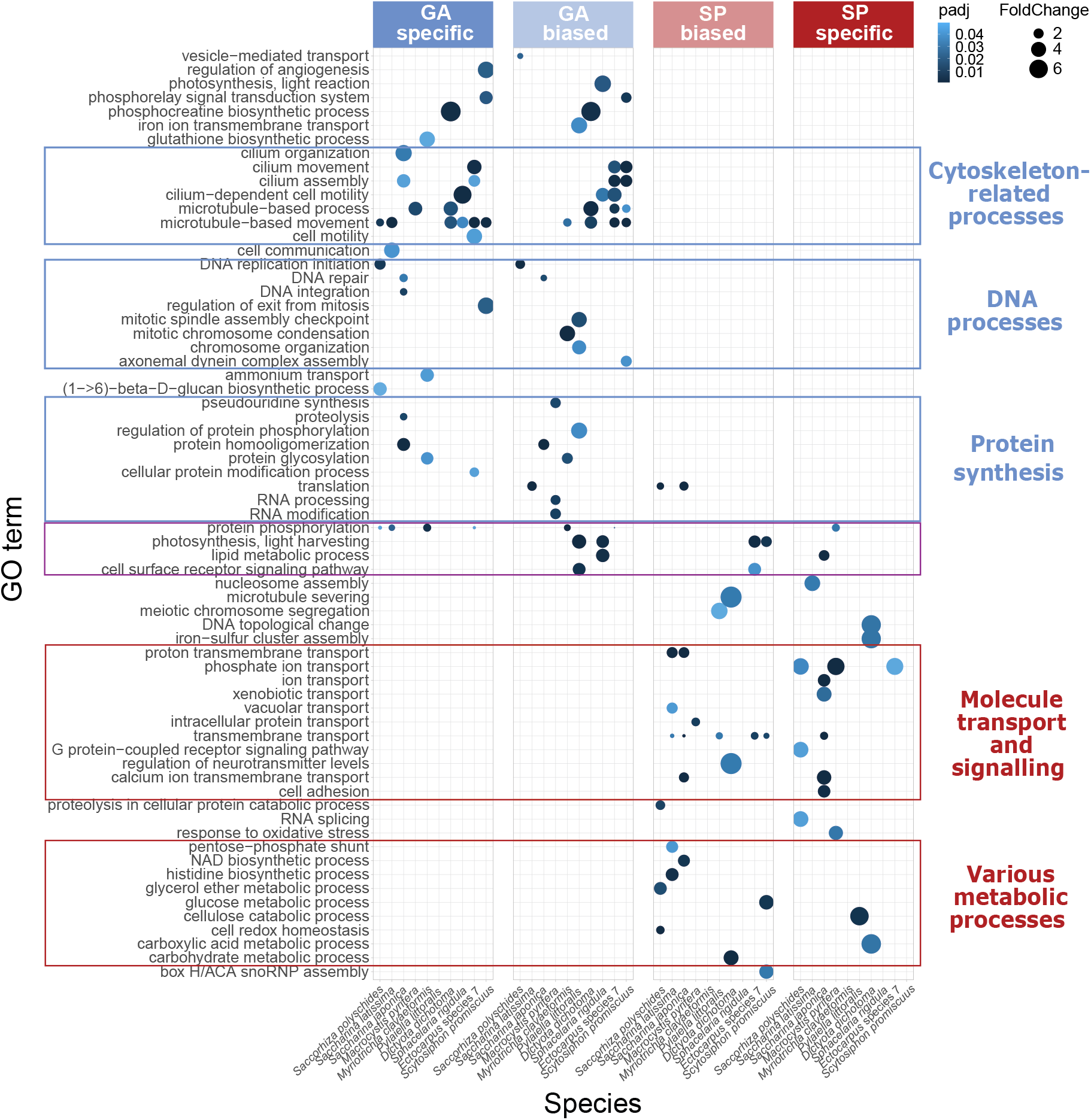
Biological function gene ontology term enrichment in the generation-biased/specific gene sets of the ten species. Conserved general functional categories are indicated by coloured boxes. padj, *p*-value adjusted for multiple testing based on the Benjamini-Hochberg false discovery rate.; GA, gametophyte; SP, sporophyte.

### Changes in gene expression patterns over the course of the *Ectocarpus* species 7 life cycle reflect the cyclic transitions between life cycle stages

The above analyses specifically provided information about gene regulation patterns underlying morphological and functional differences between the adult stages of the two generations. To obtain a broader view of gene expression during the life cycle, RNA-seq samples corresponding to multiple stages of the *Ectocarpus* species 7 life cycle were analysed: replicate samples corresponding to two very early stages (24 h and 48 h after gamete release; Figure S2) of sporophyte development were generated and combined with publicly-available samples corresponding to eight other stages of the life cycle (Table S1).

In a Principal Component Analysis (PCA) the dispersion of the samples corresponding to these ten life cycle stages mirrored the cyclic organisation of the alga life cycle (Figure 3A). Together, the two main axes explained 57% of the observed variability, with adult stages grouping together at one end of the first axis and early stages localised at the other end. The second axis approximately separated the sporophyte and gametophyte phases of the life cycle. The PCA pattern indicated gradual changes in transcriptome content between each stage of the life cycle, so that successive stages possessed overlapping but different transcriptomes.

**Figure 3.**
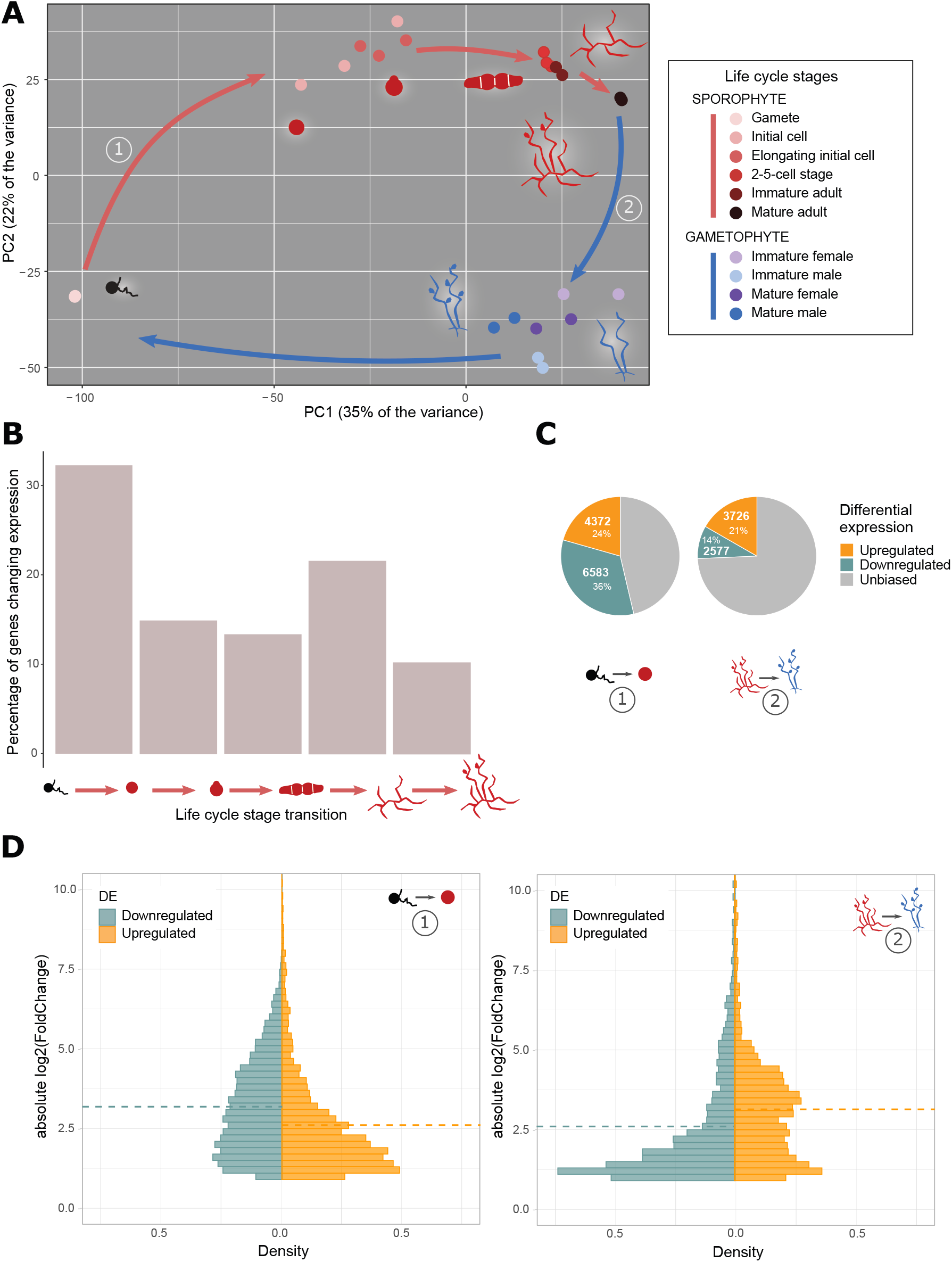
Gene expression patterns during the *Ectocarpus* species 7 life cycle. **A**. Principal component analysis of the *Ectocarpus* species 7 gene expression across ten life cycle stages. The numbers in circles refer to the differential gene expression analyses illustrated in C and D. Most of the stages analysed consisted entirely of non-flagellated cells with the exception of the gamete stage and mature sporophyte and gametophyte stages, which bear flagellated spores and gametes, respectively, in sporangia and gametangia, respectively. **B**. Percentage changes in gene expression (regression breakpoints) during *Ectocarpus* species 7 sporophyte development (from left to right, transitions between free-swimming male gamete, sporophyte initial cell (24 hours after gamete release), elongating sporophyte initial cell (48 hours after gamete release), sporophyte 2-5 cell stage, non-fertile adult sporophyte and fertile sporophyte). **C**. Proportions of differentially expressed genes in the transitions between 1) free-swimming male gamete and sporophyte initial cell stage, and 2) adult sporophyte and gametophyte stages, determined using DESeq2 (Table S1). **D**. Density distribution of the |log_2_(FoldChange)| for upregulated and downregulated genes in the differential gene expression analysis described in C. Dotted lines indicate mean values.

### The first day of sporophyte development corresponds to a major shift in the pattern of gene expression

The PCA also indicated marked differences between the gamete stage and all the other stages of the life cycle (Figure 3A). To further investigate this observation, the Trendy R package (Bacher *et al*., 2018) was used to calculate the fit of segmented regression models to individual gene expression patterns in order to identify individual genes that best matched the overall pattern of regression breakpoints. When this approach was applied to the sporophyte developmental series, a good regression fit (R^2^ > 0.5) could be calculated for 5,196 genes and 32% of these genes exhibited a change in expression (i.e. a regression breakpoint) during the transition from the gamete stage to the sporophyte initial cell (24 h) stage (Figure 3B), indicating that this transition involves a major shift in the pattern of gene expression.

To complement the above analysis, DESeq2 (Love *et al*., 2014) was used to identify genes that were differentially expressed between the free-swimming gamete and the sporophyte initial cell stage. This analysis detected 10,955 differentially expressed genes, corresponding to 4,372 and 6,583 significantly up- and down-regulated genes, respectively. These differentially expressed genes represent 58% of the total number of genes in the *Ectocarpus* species 7 genome (Figure 3C). For comparison, when adult sporophytes and gametophytes stage were compared (Figure 1B), 3,726 upregulated and 2,577 downregulated genes (35% of the total number of genes overall) were detected. The DESeq2 analysis therefore supported the conclusion that a marked change in gene expression pattern occurred between the free-swimming gamete stage and the sporophyte initial cell stage.

Interestingly, in the comparison between gametes and the sporophyte initial cell, the mean fold change in transcript abundance for downregulated genes at the transition between free-swimming gamete and initial cell was significantly higher than the mean fold change for genes that were upregulated at this transition (Wilcoxon rank sum test *p-*values < 2.10^−16^; Figure 3D). This observation suggested narrower expression ranges for the downregulated genes and this hypothesis was supported by a comparison of tau values (a measure of breadth of expression) for the downregulated and upregulated genes, which showed that the downregulated genes had significantly higher tau values (Wilcoxon rank sum test *p-*values < 2.10^−16^; Figure S3). A comparison across the ten life cycle stages also showed that 61% of the genes downregulated at the gamete to initial cell transition were most strongly expressed at the gamete stage (data in Tables S4, S5).

### Analysis of gene co-expression modules indicated coordinated regulation of genes associated with several cellular processes

To further analyse gene expression patterns over the life cycle, a weighted correlation network analysis (WGCNA) approach (Langfelder and Horvath, 2008) was used to group genes according to their patterns of expression across the ten life cycle stages. Using a cut-off of normalised counts ⌇45 (DESeq2 analysis), 16,077 of the 18,370 *Ectocarpus* species 7 genes were classed as being significantly expressed at at least one stage of the life cycle. Of these genes, 14,277 could be assigned to one of 23 gene expression modules, which varied in size from 42 to 2,414 genes (Figures S4, 4, Table S6). The modules were given arbitrary colour names (Figure 4F). The 24th “grey” module groups the 1,800 genes that were not assigned to a co-expression module (Figure 4F, Table S6).

**Figure 4.**
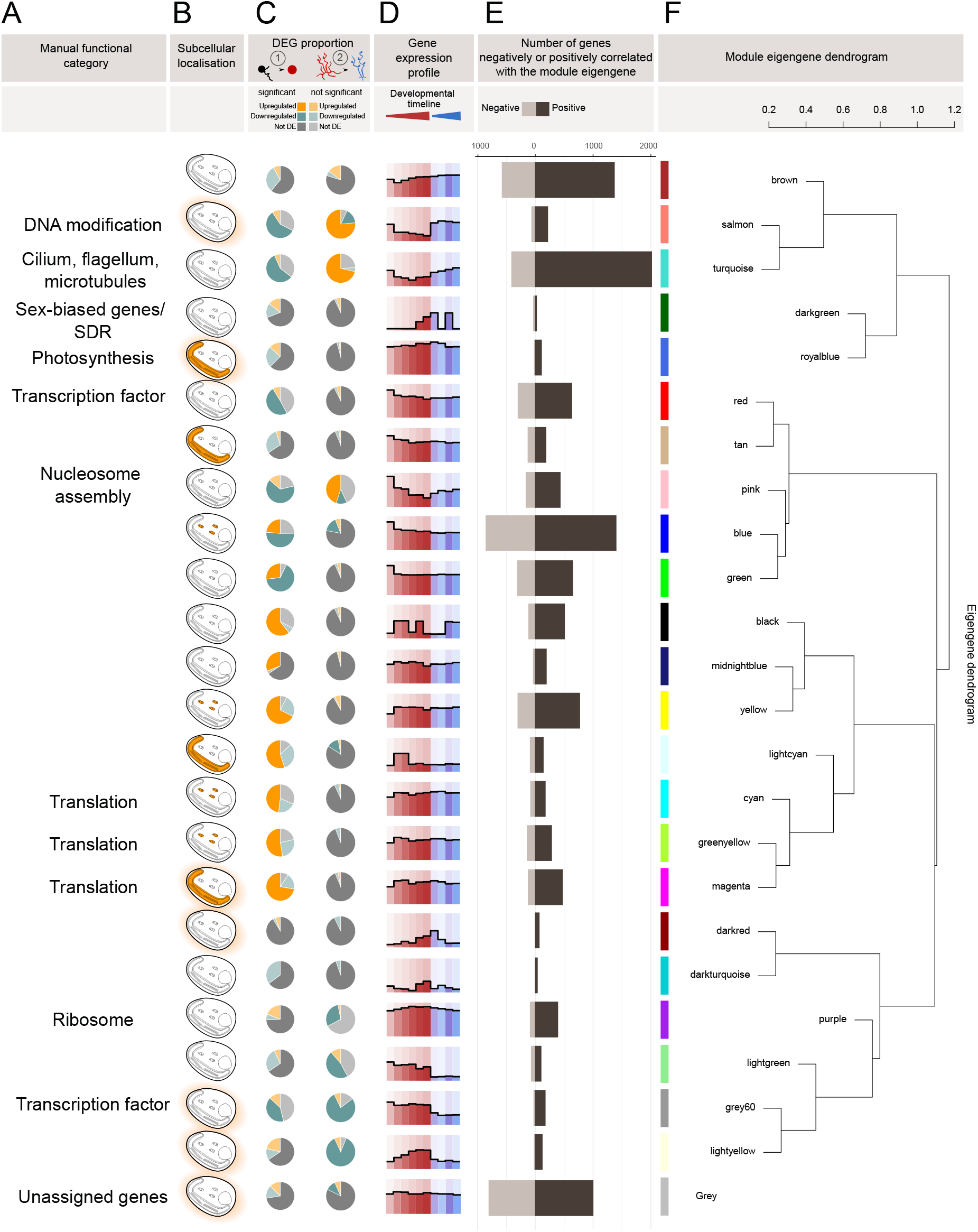
Characterisation of gene co-expression modules in *Ectocarpus* species 7. **A**. Functional categories manually assigned to modules based on Gene Ontology term enrichment analysis of each module. **B**. Enrichments in sub-cellular localisations based on HECTAR predictions: mitochondria (small ellipses), plastid (tube shape with thylakoids) and signal peptide/anchor peptide for secreted proteins. Note that HECTAR does not predict nuclear localisation. **C**. Proportion of differentially expressed genes (DESeq2 adjusted *p*-value < 0.05 and |log2FoldChange| > 1) in each gene module in the transitions between 1) free-swimming male gametes and sporophyte initial cell stage, and 2) adult sporophyte and gametophyte (Table S1). Significant enrichment (indicated by a darker shade of green or orange) means that the proportion of differentially expressed genes in the module was significantly greater (ClusterProfiler adjusted *p*-value < 0.05) than the proportion for the entire genome. **D**. Average module gene expression profile computed on genes with a WGCNA module MM > 0.86 for, from left to right: free-swimming male gamete, sporophyte initial cell (24 hours after gamete release), elongating sporophyte initial cell (48 hours after gamete release), sporophyte 2-5 cell stage, non-fertile adult sporophyte, fertile sporophyte, non-fertile female and male gametophytes, fertile female and male gametophytes developmental stages. **E**. Number of genes in the module positively or negatively correlated with the module eigengene. **F**. Module eigengene dendrogram showing the relationship between module eigengenes.

The 23 gene co-expression modules were analysed for enriched gene ontology (GO) terms (Figures 5 and S5). For 11 of the 23 gene co-expression modules, the set of enriched GO terms were sufficiently related to allow a general biological function to be manually assigned to the module (Figure 4A). These modules indicated that at least a proportion of the genes involved in several cellular processes (photosynthesis, DNA modification, flagella biosynthesis/function, transcription and translation) are co-ordinately regulated during the life cycle.

**Figure 5.**
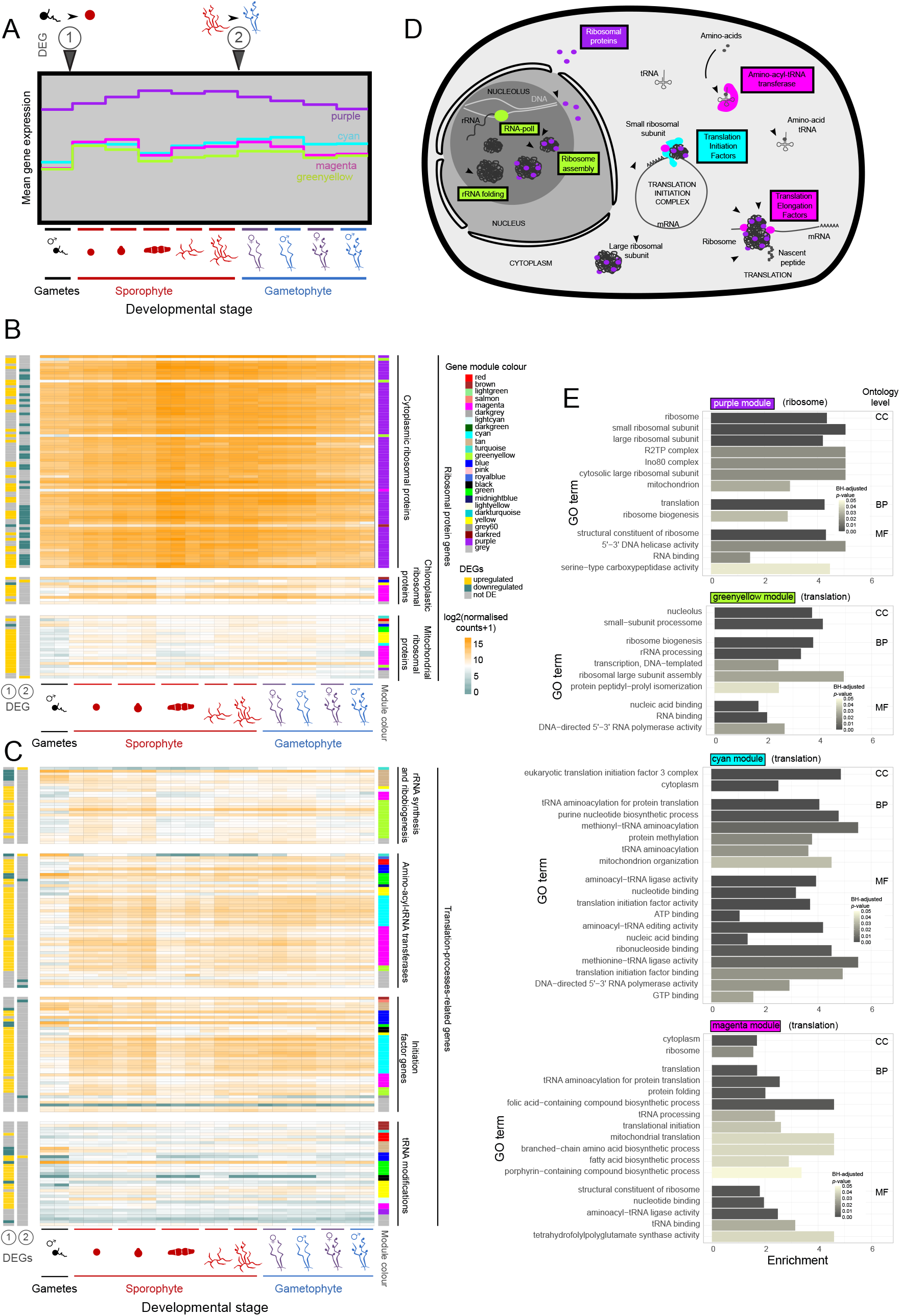
Expression of translation-related genes during *Ectocarpus* species 7 development. **A**. Mean expression profile computed for genes with a WGCNA module MM > 0.86 for (from left to right): free-swimming male gamete, sporophyte initial cell (24 hours after gamete release), elongating sporophyte initial cell (48 hours after gamete release), sporophyte 2-5 cell stage, non-fertile adult sporophyte, fertile sporophyte, non-fertile female and male gametophyte, fertile female and male gametophyte developmental stages in modules “purple”, “cyan”, “greenyellow”, “magenta”. B and C. Heatmaps showing log_2_(NormalisedCounts+1) values across the same developmental timepoints as in A for manually reannotated ribosomal protein genes corresponding to cytosolic, chloroplast and mitochondrial ribosomal subunits (B) and translation-related genes (C). Left annotation track: differential expression analysis results for the transitions between 1) free-swimming male gamete and sporophyte initial cell stage, and 2) adult sporophyte and adult gametophyte; right annotation track: WGCNA module colours. D. Schematic representation of translation-related functions enriched in the “greenyellow”, “purple”, “magenta”, “cyan” modules. When it was possible to manually assign a general function to a module, the annotation is indicated in brackets after the module name. E. Gene ontology terms significantly enriched in the sets of genes clustered within the “purple”, “greenyellow”, “cyan”, and “magenta” modules in *Ectocarpus* species 7. Enrichment is indicated as log2 of the ratio of the proportion of genes assigned to the GO term in the module divided by the proportion for the whole genome. CC, cellular component; BP, biological process; MF, molecular function.

The PCA and differential expression analyses described above provided evidence for a large-scale modification of the transcriptome during the transition between the gamete and sporophyte initial cell stage. As expected, changes in expression level at this transition strongly influenced the grouping of genes into expression modules (Figure 4F) and several modules were enriched in genes that had been identified as differentially expressed at this first transition (Figure 4C). The following two sections will focus on these modules to further characterise this important early developmental transition.

### Sporophyte down-regulated gene modules highlighted DNA- and flagella-related functions in gametes

Seven modules were significantly enriched in genes that were downregulated during the transition from gametes to the sporophyte initial cell (Figure 4C, Table S6). For three of these modules, “pink”, “salmon” and “turquoise”, the genes were also upregulated during the gametophyte stages (Figure 4D). The “pink” module was enriched in gene ontology terms related to nucleosome assembly (Figure S5) and the “salmon” module, which contained a large number of monoexonic genes, was enriched in genes predicted to have DNA-related functions (enriched gene ontology terms included “DNA integration”, “DNA recombination”, “DNA repair”, “DNA helicase” and “telomere maintenance”; Figure S5). The “turquoise” module, which had an expression profile very similar to that of the “salmon” module (Figure 4D), was enriched in genes with predicted functions related to the flagella (enriched gene ontology terms included “dynein complex”, “cilium”, “BBSome”, “microtubule-based movement”, “axonemal dynein complex assembly”, “cilium movement” and “microtubule motor activity”), and therefore presumably important for gamete motility (Figure S5). The upregulation of “turquoise” module genes in gametophytes is probably explained by the production of flagellated gametes in the gametangia of fertile thalli although it is not clear why the genes were also expressed a quite a high level in pre-fertile, adult gametophytes. It is possible that transcripts already accumulate at this stage to prepare for the production of large numbers of gametes at fertility or that some of these genes are also involved in other cellular functions, for example related to the cytoskeleton.

Cell-wall biosynthesis functions were not significantly enriched in the “turquoise”, “salmon” and “pink” modules but these modules contained the majority of annotated genes from two families of carbohydrate-active enzymes (CAZymes), the GT23 family (four genes in the “turquoise” and two genes in the “salmon” module out of ten genes in total) and the PL41 family (nine genes in the “turquoise” module, five genes in the “salmon” module and three genes in the “pink” module out of 22 expressed, annotated PL41 genes; Figure S6). The GT23 family is predicted to encode fucosyltransferases, which would transfer fucosyl groups onto chitin oligosaccharides or glycoproteins, and the PL41 family contains mannose-specific alginate lyases (Inoue and Ojima, 2019; Mazéas *et al*., 2024). While the role of GT23 enzymes in gametes is unclear, it is possible that the polysaccharide depolymerisation action of PL41 lyases may facilitate gamete release from plurilocular sporangia in a manner analogous to the actions of pectate lyases and rhamnogalacturonan lyases during fruit ripening in land plants (Méndez-Yañez *et al*., 2020; Uluisik and Seymour, 2020; Al-Hinai *et al*., 2024).

The remaining four modules that were significantly enriched in genes downregulated during the transition from the gamete to the early sporophyte (“blue”, “green”, “red” and “grey60”) did not exhibit gene upregulation during the gametophyte generation (Figure 4C, D, Table S6). No significant GO term enrichment was detected for the “blue” and “green” modules. The “red” and “grey60” modules were enriched in transcription factors and related GO terms such as “DNA-binding domain” (Figure S5). To investigate this point further a recently established dataset of 325 transcription-associated proteins (TAPs) for *Ectocarpus* species 7 (Denoeud *et al*., 2024) was enriched with the 90 Esv-1-7 domain proteins (Macaisne *et al*., 2017) from this species (Table S5, Figure S7), as the latter have been hypothesised to function as zinc finger transcription factors (Macaisne *et al*., 2017), and the expression of these genes analysed in gametes. Transcripts of approximately a third of the genes (147) in this transcription-associated dataset were more abundant in gametes than at the sporophyte initial cell stage (24 h) but the proportion was similar to the proportion for all the genes in the genome indicating that there was not an overall enrichment of the gamete transcriptome in transcription-associated genes.

Taken together, analysis of the seven modules highlighted the importance of DNA-related functions, including nucleosome assembly and transcription, and flagella-related functions in gametes.

### Subsets of translation-related genes are upregulated in the sporophyte initial cell but the gamete already expresses some translation-related and photosynthesis genes

Nine modules were significantly enriched in genes upregulated during the transition between gametes and the sporophyte initial cell (Figure 4C, Table S6). Most of the nine modules could not be clearly associated with a general cellular process but three (“cyan”, “magenta” and “greenyellow”) were enriched in functions related to translation (Figures 4A, 5). These three modules exhibited very similar expression profiles (Figure 5A) but were enriched in different classes of translation-related gene (Figure 5B-E). The “greenyellow” module contained the RNA polymerase I gene which synthesises rRNA and was enriched in GO terms related to “nucleolus”, “small-subunit processome”, “ribosome large subunit assembly”, “ribosome biogenesis” and “rRNA processing”. This module was therefore essentially related to the biosynthesis of the ribonucleic component of the ribosome (Figure 5D). The “cyan” and “magenta” modules were enriched in GO terms such as “tRNA aminoacylation for protein translation”, “tRNA processing”, “translation initiation” (Figure 5E). These two clusters included 25 of the 49 aminoacyl-tRNA-transferases annotated in the *Ectocarpus* species 7 genome, and also included a number of genes related to transcription initiation (Figure 5C). Notably, the “cyan” module included nine eIF3-subunit-encoding genes out of a total of 15 annotated genes in the genome (Figure 5C). Interestingly, the “cyan” and “greenyellow” modules were enriched (*p* <0.05 clusterProfiler) in genes with a mitochondrial targeting sequence (Figure 4B), whereas the “magenta” module was enriched (*p*<0.05 clusterProfiler) in genes that encode proteins targeted to the plastid. The plastid-targeted proteins had predicted functions other than photosynthesis (e.g. translation and transport).

The “cyan”, “greenyellow”, and “magenta” modules clustered together based on their average expression profiles (Figure 4F), suggesting that the three groups of translation-related functions share some level of co-expression. Taken together, the gene content of these three clusters suggest that there is a marked increase in translation in the initial cell compared to the gamete, both in the cytosolic and the organellar compartments. Importantly, however, not all translation-related genes exhibited marked upregulation during the transition from the gamete to the initial cell stage. For example, the transcripts of the many ribosomal-protein-encoding genes that were grouped in the “purple” module were already present in gametes at relatively high levels (Figures 5B, S8). The ribosomal protein genes are an interesting case because, whilst the majority of these genes were assigned to the “purple” module, corresponding essentially to constitutive expression, a minority of the members of this broad family exhibited more complex expression patterns and were assigned to other modules (Figures 5B, S8). In general, this latter minority of genes tend to encode proteins that are located in close vicinity in the P-stalk within the ribosome (Figure S9). The functional significance of this clustering is unclear but it has been proposed that the P-stalk influences translation fidelity and the pool of mRNAs associated with the ribosome (Wawiórka *et al*., 2017; Dopler *et al*., 2024). We also noted that ribosomal protein genes with non-constitutive expression patterns tended to have paralogues that were constitutively expressed (Figure S8). Interestingly, cytosolic and organellar ribosomal protein genes did not exhibit the same patterns of expression with respect to the life cycle, with the genes encoding organellar proteins exhibiting more complex expression patterns (i.e. they were assigned to multiple modules; Figure 5B).

The “royalblue” module is another example of a module which did not exhibit marked changes in expression between the gamete and the sporophyte initial cell. This module was highly enriched in genes related to photosynthesis or encoding chloroplast-targeted proteins (Figure 4A,B,C). Based on the transcriptomic data, therefore, both cytosolic translation and photosynthesis appear to be processes that are already important at the gamete stage, although it is possible that transcripts related to these processes may accumulate in gametes but be translated at a later stage.

### Comparative analysis of life-cycle-related gene expression between *Ectocarpus* species 7 and *D. dichotoma*

To determine whether the developmental and life-cycle-related transcription patterns observed in *Ectocarpus* species 7 corresponded to common, fundamental features that are shared with other brown algae, a comparative analysis was carried out between *Ectocarpus* species 7 and *D. dichotoma*. These two species are phylogenetically distant within the brown algal tree (Akita *et al*., 2022). Based on a recent estimate (Choi *et al*., 2024), their common ancestor dates to about 250 Mya. Consequently, if a feature is shared between the two species, this is an indication that it may have been conserved over a significant part of the evolutionary history of the brown algal lineage.

Replicated transcriptomic data was available for seven stages of the *D. dichotoma* life cycle (Table S1). In a PCA analysis (Figure S10) the adult sporophyte and gametophyte stages clustered together, presumably reflecting the isomorphy between the two generations. Consequently, the distribution of stages in the PCA did not follow the progression of the life cycle, as had been observed for *Ectocarpus* species 7. However, the first dimension clearly separated male gametes from the other *D. dichotoma* life cycle stages and the second dimension separated early developmental stages from adult stages. *D. dichotoma* has flagellated male gametes and non-flagellated egg cells. The latter were clustered with early sporophyte stages (zygote and 8h embryo), suggesting that the markedly different gene expression patterns observed in male gametes from the two species compared to other life cycle stages were related to their being actively swimming, flagellated cells rather than being a universal characteristic of all brown algal gametes.

Life-cycle-related gene expression in *Ectocarpus* species 7 and *D. dichotoma* was compared using two different approaches. The first aimed to determine whether the co-expression patterns of *Ectocarpus* species 7 genes grouped into modules were mirrored by the *D. dichotoma* orthologues of these genes. The second involved independently clustering *D. dichotoma* genes into co-expression modules using life cycle transcriptomic data and then determining whether there were similarities between the modules established for the two species.

To optimally identify conserved one-to-one orthologues between *Ectocarpus* species 7 and *D. dichotoma*, an orthologue analysis was carried out specifically for the two species using Orthofinder (Emms and Kelly, 2019). This approach identified 8,527 one-to-one orthologous relationships between the 18,370 *Ectocarpus* species 7 genes and the 20,583 *D. dichotoma* genes (Table S7). This orthologue dataset was used to compare gene expression patterns in the two species.

### *D. dichotoma* orthologues of sets of genes from *Ectocarpus* species 7 modules exhibited conserved co-expression for some modules that grouped genes involved in the same cellular process

To determine whether the co-expression patterns of genes in *Ectocarpus* species 7 modules was conserved in *D. dichotoma*, the *Ectocarpus* species 7 genes in each module were substituted with their *D. dichotoma* orthologues and various parameters, notably the preservation of the density of the module and the preservation of the connectivity, were evaluated (Table S8) using the *D. dichotoma* transcriptomic data for the seven different life cycle stages (Table S1). The density preservation of a module was measured by evaluating whether the average correlation for the module was conserved in the second species and whether the module summary profile (module eigengene) still explained a high proportion of the variance after the genes were substituted with orthologues from the second species. Connectivity preservation, on the other hand, determined whether the correlation pattern between the genes was conserved after substitution. Various density and connectivity statistics based on functions in both the WGCNA R package and the NetRep R package were aggregated to estimate conservation of gene co-expression for each *Ectocarpus* species 7 module in *D. dichotoma*.

Generally, the modules with the best conservation statistics in this analysis were those that had been identified as being highly enriched in *Ectocarpus* species 7 genes with predicted housekeeping functions (Figure 6A). Thus, the “photosynthesis” (“royalblue”), the “ribosome” (“purple”), the “cilium and flagellum” (“turquoise”) and various translation-associated (“greenyellow”, “cyan”, and “magenta”) modules all showed a high level of conservation. The expression pattern of the “darkgreen” module was also considered to be conserved in *D. dichotoma* despite the fact only three orthologues were detected. The “darkgreen” module consists principally of genes located within the non-recombining sex-determining region (SDR) of the female (U) and male (V) sex chromosomes (Ahmed *et al*., 2014) and is therefore a special case because the co-expression pattern is determined not only by gene expression but also by gene presence (i.e. presence of the U or V sex chromosome in gametophytes). The module was considered to be conserved in *D. dichotoma* because the three orthologues are located in the SDR in both species.

**Figure 6.**
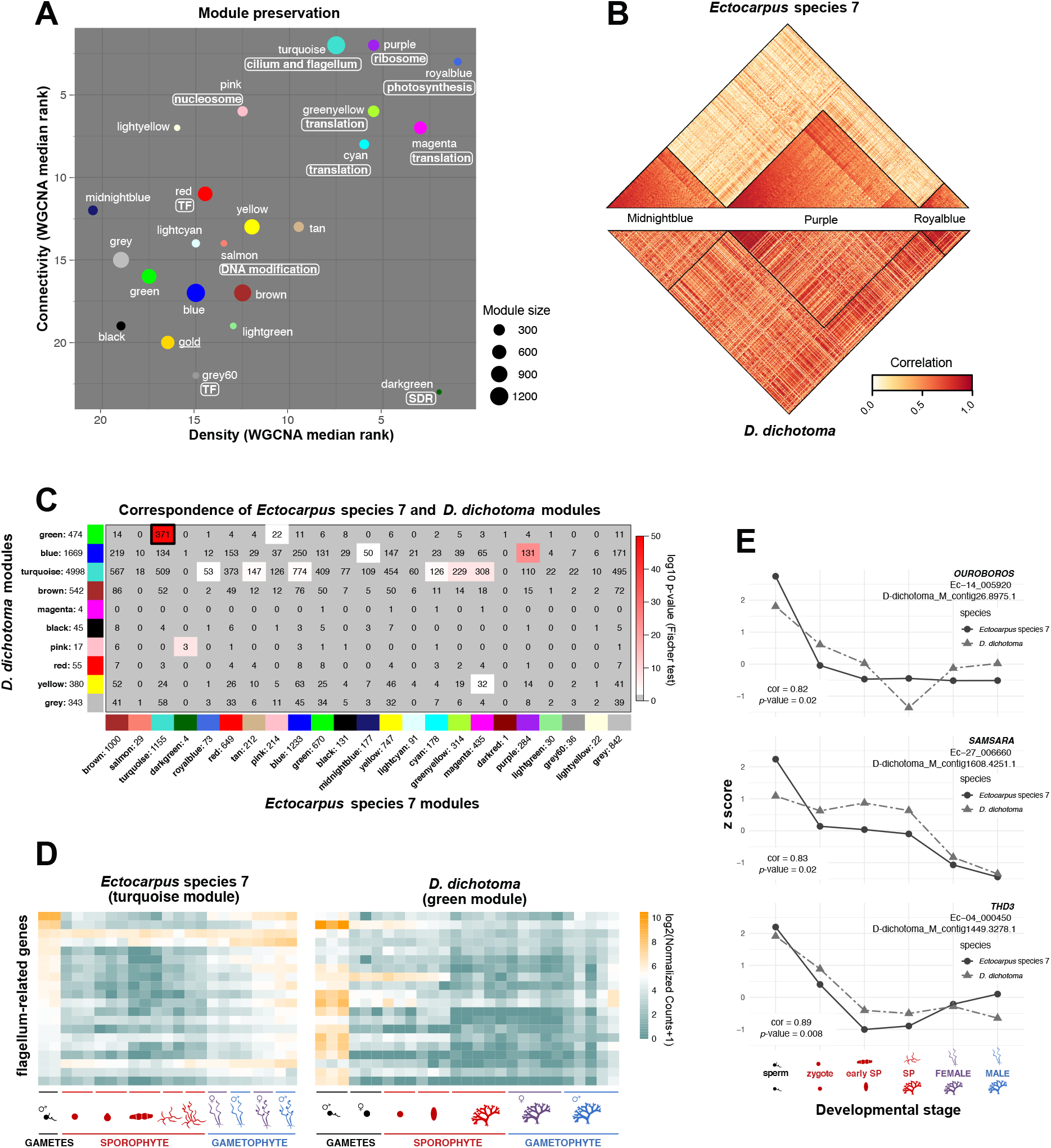
Conservation of life-cycle-related gene co-expression patterns between *Ectocarpus* species 7 and *D. dichotoma*. **A**. WGCNA density and connectivity median rank statistics indicating the degree of conservation of gene co-expression patterns when of *Ectocarpus* species 7 genes in *Ectocarpus* species 7 co-expression modules were substituted with their *D. dichotoma* orthologues. Manually-assigned general biological functions are indicated in boxes. SDR, sex-determining region. Module size indicates the number of one-to-one orthologues in each module. The modules “darkred” and “darkturquoise” were not included in this analysis because they had only one and zero orthologues, respectively (Table S6). TF, transcription factor; SDR, sex-determining region. **B**. Correlation heatmap comparing three *Ectocarpus* species 7 modules between *Ectocarpus* species 7 and *D. dichotoma*: “midnightblue” (very poor conservation), “purple”, and “royalblue” (good conservation). The lower half of the heatmap was calculated based on the expression pattern of the *D. dichotoma* orthologue of each *Ectocarpus* species 7 gene in each module. C. Counts of shared one-to-one orthologues between gene co-expression modules defined for *Ectocarpus* species 7 (x-axis) and *D. dichotoma* (y-axis). The colour code represents the log_10_(*p*-value) for Fisher’s exact test (red bar), which was applied to determine whether pairs of modules shared a greater number of one-to-one orthologues than expected from a random distribution. Numbers after the module names indicate the number of one-to-one orthologues in each module. The “darkturquoise” module was not included in this analysis because it contained zero orthologues (Table S6) D. Heatmap showing the expression levels (log_2_(NormalisedCounts+1)) of 20 selected one-to-one orthologous flagellum-related genes in *Ectocarpus* species 7 and *D. dichotoma. Ectocarpus* species 7 timepoints: free-swimming male gamete, sporophyte initial cell (24 hours after gamete release), elongating sporophyte initial cell (48 hours after gamete release), sporophyte 2-5 cell stage, non-fertile adult sporophyte, fertile sporophyte, non-fertile female and male gametophytes, fertile female and male gametophytes. *D. dichotoma* timepoints: sperm, egg cell, zygote, embryo, adult non-fertile sporophytes, and female and male gametophytes. **E**. Computed z-score based on log_2_(NormalisedCounts+1) values for three TALE homeodomain transcription factors over six developmental stages shared between *Ectocarpus* species 7 and *D. dichotoma*, namely sperm, zygote, early sporophyte, non-fertile sporophyte, female and male gametophyte. The Pearson correlation coefficient between the two expression datasets (cor) and the associated *p*-value are indicated.

For modules that had good conservation statistics between the two species, *D. dichotoma* orthologues of the *Ectocarpus* species 7 genes that were best correlated with the module eigengene also exhibited a strong correlation with the module eigengene after the *Ectocarpus species* 7 genes had been replaced with the *D. dichotoma* orthologues (Figure 6B). For example, for the “purple” and “royalblue” modules (which had good conservation statistics; Figure 6A) in the top part of Figure 6B, the *Ectocarpus* species 7 genes that were best correlated with the module eigengene are positioned on the left. In the bottom part of the figure the *D. dichotoma* orthologues of these genes are also positioned on the left and they also show strong correlation with the module eigengene. This conservation of correlation with the module eigengene between orthologues was not observed for modules such as “midnightblue” that had poor conservation statistics between the two species (Figure 6B).

To summarise, this analysis indicated that the co-expression pattern of the genes in a *Ectocarpus* species 7 module following replacement with the *D. dichotoma* orthologues was only conserved for a subset of the modules. The modules that exhibited conservation of co-expression across the two species in this analysis tended to contain sets of genes that functioned in a basic, evolutionarily-conserved cellular process. Moreover, conservation of module co-expression patterns across the two species appeared to be conferred preferentially by orthologues of the *Ectocarpus* species 7 genes that showed the best correlation with the module eigengene.

### A subset of life-cycle-related gene co-expression modules were conserved between *Ectocarpus* species 7 and *D. dichotoma*

For the second approach to compare life-cycle-regulated gene expression in *Ectocarpus* species 7 and *D. dichotoma*, WGCNA was used to define co-expressed gene modules independently for *D. dichotoma* using the replicated transcriptomic data for the seven different *D. dichotoma* life cycle stages (Table S1; Figure S4B). The number of modules defined for *D. dichotoma* was lower than the number that had been obtained for *Ectocarpus* species 7 (nine compared with 23; Figure S10, Tables S6, S9, S10), presumably because data was available for fewer life cycle stages and because the *D. dichotoma* adult sporophyte and gametophyte generations are isomorphic and therefore have more similar overall gene expression patterns (Figure S10A). As for the *Ectocarpus* species 7 modules, analysis of enriched GO terms (Figure S11) allowed general biological functions to be manually assigned to a subset of the modules (five of the nine modules; Figure S10A).

Correlation between modules across the two species was assessed based on the number of shared one-to-one orthologues in pairwise comparisons. Fisher’s exact test was used to determine whether *Ectocarpus* species 7 and *D. dichotoma* modules shared a greater number of one-to-one orthologues than expected from a random distribution. This analysis identified several pairs of life-cycle-related gene co-expression modules with significantly conserved gene contents between the two species but not all modules were conserved (Figure 6C). As in the first analysis described above, conservation tended to be strongest for *Ectocarpus* species 7 modules that had been manually assigned general functional annotations and which had registered the highest values for module preservation statistics in the first analysis (Figure 6A).

“Purple” was one of the most strongly conserved *Ectocarpus* species 7 modules: 131 of its 284 one-to-one orthologues were shared with the *D. dichotoma* “blue” module (Figure 6C), with most of the shared orthologues predicted to encode ribosomal proteins. However, the *D. dichotoma* “blue” module exhibited a marked downregulation in male gametes, which was not observed for the *Ectocarpus* species 7 “purple” module. In contrast, the *Ectocarpus* species 7 “turquoise” module, shared 371 (of a total of 1155) one-to-one orthologues with the *D. dichotoma* “green” module, principally with predicted cilium- and flagellum-related functions (Figure 6C,D), and these two modules also exhibited similar life-cycle-related expression patterns, in particular high levels of expression in male gametes. Therefore, both the gene content and the expression patterns of these two modules appear to have been conserved over evolution. Most of the one-to-one orthologues in the translation-related *Ectocarpus* species 7 modules “cyan”, “greenyellow” and “magenta” corresponded with genes in the *D. dichotoma* “turquoise” module, consistent with the poorer resolution of the *D. dichotoma* modules (Figure 6C). Nonetheless, all of these modules exhibited a lower level of gene expression in male gametes, indicating conservation of their expression patterns over evolutionary time.

Taken together, the comparison between *Ectocarpus* species 7 and *D. dichotoma* indicated that there has been partial conservation of life-cycle-related gene expression patterns between the two species, in particular for modules corresponding to clearly-identifiable cellular functions. Moreover, some modules exhibited conservation not only in terms of gene (orthologue) content but also exhibited similar expression patterns over the course of the life cycle, proving further support for the (partial) conservation of life-cycle-related gene expression patterns over time.

The above analyses provided no evidence that transcription-related modules were conserved, overall, between *Ectocarpus* species 7 and *D. dichotoma* but a more detailed analysis was carried out to determine whether life-cycle-related expression patterns might have been conserved for individual transcription factor genes. For this, 214 transcription factors with one-to-one orthologues in the two species were selected and their expression patterns (normalized expression z-scores) analysed across six life cycle stages that were approximately equivalent between the two species (gamete, zygote, early sporophyte, adult sporophyte, adult male gametophyte and adult female gametophyte; Figure S12, Table S1). High Pearson correlation coefficients were obtained for a subset of the transcription factors but none of the correlations were significant at *p* = 0.05 after correction for multiple testing (Table S11). However, the applied correction was stringent because a large number of orthologous genes were considered so the most strongly correlated transcription factors may nonetheless merit further study in the future. We noted, for example, that all three members of the TALE homeodomain transcription factor family, which includes the life cycle regulators *ORO* and *SAM* (Arun *et al*., 2019), were among the genes that exhibited high Pearson correlation coefficients in this analysis (Table S11, Figure 6E).

## Discussion

### Subsets of genes with related functions are co-expressed during the life cycle

This study analysed replicated RNA-seq samples for ten different stages of the *Ectocarpus* species 7 life cycle to provide an overview of life-cycle-related gene expression in this species. Clustering of genes into co-expression modules indicated that subsets of the genes associated with several cellular processes, including transcription, protein synthesis, flagella biosynthesis/function and photosynthesis, are co-ordinately regulated during the life cycle.

The coordinated expression patterns of the genes in these modules implies coordinated regulation of transcription and/or mRNA stability over the course of the life cycle. The regulatory systems that mediate this coordinated expression are currently unknown for brown algae but information about the mechanisms that co-ordinately regulate these processes in other eukaryote lineages can provide some indications of possible regulators. For example, in *Ectocarpus* species 7, four different modules were enriched in translation-related genes. In other eukaryotic lineages, a large proportion of ribosomal protein genes are regulated by the mTOR/PKA pathway (Martin *et al*., 2004; Ni and Buszczak, 2023). Interestingly, genes predicted to encode cytosolic and organellar ribosomal proteins were regulated differently in *Ectocarpus* species 7. Note also that this conservation of expression patterns provides additional support for the transfer of the *Arabidopsis* ribosomal protein gene nomenclature (Scarpin *et al*., 2023) to *Ectocarpus* species 7 genes (https://bioinformatics.psb.ugent.be/orcae/overview/EctsiV2) based on sequence similarity.

In land plants, photosynthesis-related genes have been shown to be regulated by both light and developmental cues and the regulatory pathways have been characterised. Phytochrome signalling counteracts negative regulation of photosynthesis genes by transcription factors such as PHYTOCHROME-INTERACTING FACTOR (PIF), PIF-LIKE (PIL) and GOLDEN2-LIKE1 (GLK1), which act downstream of various developmentally regulated pathways (Waters *et al*., 2009; Wang *et al*., 2017). However, orthologues of these transcription factors have not been found in brown algae and, consequently, coordinated regulation of photosynthetic genes in brown algae, as was observed here for the “royalblue” module, presumably involves a different set of actors.

As far as flagella- and cilium-related genes are concerned, these organelles are widespread in eukaryotes and the last eukaryotic common ancestor (LECA) probably possessed cilium-related structures (Carvalho-Santos *et al*., 2011). In Metazoans, two transcription factor families, FOXJ1 and RFX, control the formation of core components of all types of cilia (Choksi *et al*., 2014). However, although *Saccharomyces cerevisiae* also possesses an *RFX* gene, this family of transcription factors has only been shown to regulate ciliogenesis in metazoans and is not found outside the opisthokonts (Chu *et al*., 2010). In *Chlamydomonas reinhardtii*, ciliogenesis is under the control of XAP5 (Li *et al*., 2018), a transcription factor that is widely distributed across eukaryotic lineages. Neither *FOXJ1* nor *RFX* have been found in brown algal genomes, but *Ectocarpus* species 7 does possess an *XAP5* homologue (LocusID Ec-05_002220). This gene therefore represents a possible candidate for future analysis of the regulation of this pathway.

These examples illustrate how candidate regulatory genes may be identifiable for cellular processes that are highly conserved across eukaryotes but candidates are more difficult to identify for modules that group brown-algal-specific genes of unknown function or functions such as the PL41 and GT23 gene families that do not have equivalents in other eukaryotic lineages. In some instances, regulatory genes may show similar expression patterns to the genes they regulate as a result of auto-regulation, and therefore be included within a co-expression module, but this will not be the case for all regulators.

### Identification of marked transcriptome changes during early sporophyte development

Analysis of RNA-seq data for multiple developmental stages of *Ectocarpus* species 7 provided a broad overview of gene expression throughout the life cycle, with a particular focus on the early stages of sporophyte development. Progression through the life cycle was correlated with concomitant, progressive changes in the composition of the transcriptome (Figure 3A) and a particularly marked change in composition occurred at the transition from the gamete stage to the initial stage of sporophyte development. Most of the gene co-expression modules identified in this study exhibited changes in transcript abundance at this transition and more than half of the genes in the genome were differentially expressed between the two stages.

Periods of large-scale changes in transcriptome composition have been observed during early development of both animals and land plants (Xue *et al*., 2013; Anderson *et al*., 2017; Vastenhouw *et al*., 2019; Zhao *et al*., 2019), where they tend to coincide with the maternal-to-zygotic transition (MZT). The MZT corresponds to the switch from reliance on maternal transcripts supplied to the transcriptionally-inactive egg cell to mRNAs produced by transcription of the zygote genome (Tadros and Lipshitz, 2009; Vastenhouw *et al*., 2019). During the MZT, induction of the expression of genes in the zygote genome, which is referred to as zygotic gene activation (ZGA), coincides with a phase of degradation of maternal transcripts in both animals and land plants. The exact timing of MZT varies across species, occurring rapidly in land plants, only hours after zygote formation (Anderson *et al*., 2017; Chen *et al*., 2017; Zhao *et al*., 2019), but considerably later in animals, for example occurring three days after fertilisation in 8-cell human embryos (Xue *et al*., 2013). However, even in animals, 1-cell stage embryos exhibit a distinct transcriptome pattern (Xue *et al*., 2013).

*Ectocarpus* is not oogamous and both male and female gametes are small, flagellated cells that swim actively for a significant part of their lifetime. It therefore seems likely that these cells are already transcriptionally active during the gamete stage, in which case they would not rely solely on maternally-produced transcripts (although this has not yet been investigated experimentally). Consequently, it is not clear whether there is a specific phase of degradation of maternal transcripts in gametes equivalent to the phases of maternal transcript degradation that have been observed at the MZT in land plants and animals. However, the marked change in transcriptome composition during early development is a feature that is shared with animals and land plants, underlining the broad importance of large-scale transcriptional changes during early development.

In terms of functional categories, fewer genes involved in flagellar function or transcription were expressed after the transition from the gamete stage to the initial stage of sporophyte development. Downregulation of flagellar genes is expected for this transition from a flagellated to a non-flagellated cell. The reduction in the number of expressed transcription factor genes is more surprising but it is likely that the nature of the transcription factor genes expressed or repressed at this stage is more important than the overall number of transcription factor genes expressed. Trends in transcription factor gene up- and down-regulation are not consistent across species, with a decrease in the number of different transcription factor gene transcripts observed in *Arabidopsis* (Zhao *et al*., 2019) but an increase in rice (Anderson *et al*., 2017), maize (Chen *et al*., 2017) and human (Xue *et al*., 2013), coinciding with the MZT. Moreover, in rice and maize, specific loci that play key roles during embryogenesis such as members of the BABYBOOM and WUSCHEL gene families are upregulated at the MZT/ZGA, supporting the idea that the identity of the specific transcription factor genes expressed is more important than the overall numbers (Anderson *et al*., 2017; Chen *et al*., 2017). Note that *Ectocarpus* has AP2 domain and homeodomain proteins but no close homologues of the BABYBOOM and WUSCHEL families so it is likely that different transcriptional regulators are involved in regulating development at the equivalent stage in brown algae.

Analysis of modules that exhibited overall gene upregulation between the gamete and early sporophyte stages identified translation as a general function that was upregulated during this transition. However, the upregulation of these genes does not appear to have corresponded to *de novo* induction of translation because a subset of translation-related genes was already expressed at the gamete stage. The overall gene expression pattern is more indicative of a step increase in translation capacity or complexity in the early sporophyte compared to the gamete.

### Conservation of life-cycle-related gene expression patterns across brown algal species

Comparison of generation-biased gene expression between adult gametophyte and sporophyte stages across ten brown algal species identified a strong positive correlation between the degree of dimorphism between generations and the number of GBGs identified per species, indicating that the majority of GBG expression is related directly or indirectly to thallus morphology, at least for the adult stages analysed. Orthologue analysis indicated only weak overlap between GBG sets from different species, supporting an earlier conclusion that there is rapid turnover of GBG sets in brown algae (Lipinska *et al*., 2019). Interestingly, however, analysis of GO enrichment in GBG sets indicated that several broad functional categories were conserved across gametophyte-biased gene sets, whereas different broad functional categories were conserved across sporophyte-biased gene sets (Figure 2). This observation suggests that the two generations have conserved life-cycle-related functional characteristics that are independent of morphological complexity or the degree of dimorphism between generations.

The analysis of whole adult individuals to study GBG expression therefore identified some interesting conserved life-cycle-related features but the resolution of this approach is limited because only one sporophyte stage and one gametophyte stage was analysed per species and because, at the adult stage, the data represent an average view of the transcriptomes of multiple different cell types. The comparative analysis of multiple stages of early sporophyte development in *Ectocarpus* species 7 and *D. dichotoma* allowed a more detailed analysis of developmentally regulated gene expression and, notably, showed that there was considerable conservation of co-expressed gene modules between two species, particularly for modules that were enriched in genes involved in evolutionarily-conserved functional categories. Moreover, when the known life cycle regulatory genes *SAM* was analysed, there was clear evidence of conservation of its expression patterns across the two species. These results underline the importance of carrying out detailed transcriptomic analyses to identify and investigate conserved life cycle regulatory pathways.

## Materials and Methods

### Life cycle stages and variants on the sexual cycle

Both *Ectocarpus* species 7 and *D. dichotoma* have haploid-diploid life cycles that involve alternation between a sporophyte and a gametophyte generation (Müller, 1967; van den Hoek *et al*., 1995). During the sexual life cycle, the diploid sporophyte generation produces haploid meio-spores through meiosis. These spores germinate into haploid gametophytes that can be male or female (dioicy). Male and female gametes, produced, respectively, by male and female gametophytes, fuse to produce a diploid zygote, which, in turn, develops into the next sporophyte generation. The zoids (gametes and spores) are the only flagellated stages of the life cycle. In *Ectocarpus* species 7, the germinating zygote undergoes a developmentally symmetrical first division to produce a basal filament, which then branches to establish a basal filament system consisting of round and elongated cells. Once this basal system is established, upright filaments consisting of cylindrical cells grow up into the medium and these filaments bear the sexual structures, plurilocular sporangia and unilocular sporangia (Peters *et al*., 2008). *Ectocarpus* species 7 also exhibits variations on this basic sexual life cycle, at least in culture (Müller, 1967). For example, parthenogenesis of unfertilised gametes gives rise to haploid partheno-sporophytes. A few hours after release from the plurilocular sporangia, unfertilized free-swimming male gametes attach to the substrate, lose their flagella and acquire a round shape. Upon attachment they start to develop a cell wall. After about two days, the initial cell elongates and divides, producing the first cells of the sporophytic basal filaments. All but one of the sporophyte stages analysed in this study corresponded to partheno-sporophyte samples. Partheno-sporophytes exhibit identical morphology and reproductive functionality to diploid sporophytes and pure samples of early-stage diploid sporophytes are difficult to obtain because of a high level of gamete parthenogenesis in genetic crosses (Coelho *et al*., 2012a).

### Publicly available RNA-seq datasets

RNA-seq data for the sporophyte and gametophyte generations of the following ten species was recovered from the Phaeoexplorer website (https://phaeoexplorer.sb-roscoff.fr/) or from public databases: *Macrocystis pyrifera, Saccharina latissima, Saccharina japonica, Saccorhiza polyschides, Myriotrichia claveaformis, Pylaiella littoralis, D. dichotoma, Sphacelaria rigidula, Ectocarpus* species 7 and *Scytosiphon promiscuus*. In addition, RNA-seq data was collected for multiple stages of the *D. dichotoma* and *Ectocarpus* species 7 life cycles. All the accession numbers for these data are listed in Table S1.

### Comparisons of orthologues across multiple brown algal species

To compare orthologues across the ten brown algal species analysed for generation-biased gene expression an orthogroup analysis was carried out using Orthofinder (Emms and Kelly, 2019) version 2.5.2. Strict selection of only one-to one orthologues did not identify sufficient orthologues (1,038), principally due to gene model fragmentation in the lower quality genome assemblies. A more relaxed selection approach was therefore used, taking into account genome assembly quality. This relaxed approach, which required exactly one orthologue from the best quality assemblies (*D. dichotoma, Ectocarpus* species 7, *P. littoralis, S. latissima, S. promiscuus* and *S. rigidula*), between zero and two orthologues for *M. pyrifera*, zero or one orthologue for *S. japonica*, and any number of orthologues for *S. polyschides* and *M. clavaeformis*, identified 9,317 orthogroups (Table S3). The assumption underlying this approach was that the multiple gene models in the poor-quality genomes corresponded to fragments of a single gene but note that we cannot rule out the possibility that a minority of these groups of genes corresponded to tandem gene duplications. To calculate generation bias when a species possessed more than one gene model in a particular orthogroup, each model was given a score of +1, -1 or 0 if the expression data indicated that they were sporophyte-biased/specific, gametophyte-biased/specific or neither, respectively. The mean of these values was then calculated and the orthologue was classed as sporophyte-biased, gametophyte-biased or unbiased if the value was >0, <0 or 0, respectively.

### Algal strains and culture conditions

*Ectocarpus* species 7 strain Ec32 gametophytes were grown in sterile Provasoli-enriched (0.5x) natural sea water at 13°C under 12h:12h day:night conditions until they reached maturity. Synchronous gamete release was induced by removing excess seawater and incubating in the dark and in the cold for 1 h. Sterile Provasoli-enriched (0.5x) natural sea water (Starr and Zeikus, 1993; Coelho *et al*., 2012b) was then added and the gametophytes incubated for 30 min to allow gamete release. The medium was filtered through 1 µm cell strainer (pluriSelect, Leipzig, Germany) to eliminate any contaminating gametophytic fragments. The gamete suspension was then diluted to a concentration of 1.5 million cells per mL (A_450nm_ = 0.01) to produce a density of 300 cells per mm^2^ on plastic 150 cm Petri dishes after gamete settlement. After gamete release, the Petri dishes were incubated at 13°C under 12h:12h day:night conditions at a light intensity of 40 µmol.m^-2^.s^-1^ for 24 or 48 h. Cultures were sampled in the late afternoon to ensure homogeneity with regard to circadian rhythms. Sampling was carried out by scraping the bottom of the culture plate with a cell scraper (Sarstedt, Nümbrecht, Germany) after the seawater medium had been removed. The cell suspension was centrifuged at 14000 g for 15 min and the pellet flash-frozen in liquid nitrogen.

### RNA extraction and RNA-seq sequencing and mapping

RNA was extracted using the Macherey Nagel Nucleospin Plant and Fungi kit (Hoerdt, France). Frozen pellets were resuspended in lysis buffer (500 µL of PFL buffer, 40 µL of PFR buffer and 50 µL of 1 M DTT per sample) and the cells were lysed by three 30-second treatments at 7800 rpm in 2 mL “tough micro-organism” lysis tubes (VK05) in a Precellys Evolution homogeniser (Bertin Technologies, Montigny-le-Bretonneux, France), followed by a 5 min incubation at 56°C. RNA extraction followed the manufacturer’s instructions except that a DNA digestion step was added after the first wash (digestion on the column with 5 µL of Qiagen DNAse I in 75 µL RDD buffer for 20 s before a second wash with PFW I buffer). The PFW II buffer wash step was carried out three times. RNA was eluted in 50 µL of RNAse-free H_2_O. RNA-seq libraries were constructed after oligo-dT selection and sequenced by Genewiz (Leipzig, Germany) on an Illumina NovaSeq 6000, using a standard unstranded paired-end protocol to produce a minimum of 50 million reads per condition. Data quality was assessed with FastQC (Andrews, 2016) version 0.11.0 or 0.11.9 and reads were trimmed with Trim Galore (Krueger, 2015) version 0.6.5 with parameters --illumina --length 50 --quality 24 --stringency 6 --max_n 3. Genomes were indexed and data was mapped onto the v2 version of the *Ectocarpus* species 7 reference genome (Sterck *et al*., 2012; Cormier *et al*., 2017) using HISAT2 (Kim *et al*., 2019) version 2.1.0 with default parameters (the parameter --rna-strandedness was specified according to the information in Table S1). The output was converted to bam format and ordered and indexed using the SAMtools (Danecek *et al*., 2021) version 1.10 functions order and index. Quantification of transcript abundance was carried out using the featureCounts function (Liao *et al*., 2014) from the Subread package version 2.0.1 on CDS features grouped by Parent or LocusID.

### Identification of differentially expressed genes

Differentially expressed genes were identified using DESeq2 (Love *et al*., 2014) version 1.34.0 in R version 4.1.2. Genes were considered to be differentially expressed if they had an adjusted (Benjamini-Hochberg false discovery rate) *p*-value of <0.05 and an absolute log_2_(FoldChange) greater than 1. Breakpoints in gene expression trends were analysed using the Trendy R package (Bacher *et al*., 2018) version 1.16.0 with the functions trendy (maxK = 3, pvalCut = 0.05) and topTrendy (adjR2Cut = 0.5). DESeq2 version 1.34.0 was also used to calculate normalised counts (NormalisedCounts) using the median of ratios method. Transcripts per kilobase million (TPM) values were calculated from counts of mapped reads using an in-house custom script. Based on the differential expression analyses and the TPM values, genes were classified into six categories: gametophyte-biased (mean gametophyte and mean sporophyte TPM values both ⍰1, log2(fold change) ⍰1, adjusted *p*-value <0.05), sporophyte-biased (mean gametophyte and mean sporophyte TPM values both ⍰1, log2(fold change) ⍰-1, adjusted *p*-value <0.05), gametophyte-specific (mean TPM < 1 for the sporophyte and ⍰1 for the gametophyte, log2(fold change) ⍰1, adjusted *p*-value <0.05), sporophyte-specific (mean TPM <1 for the gametophyte and ⍰1 for the sporophyte, log2(fold change) ⍰-1, adjusted *p*-value <0.05), unbiased genes (mean gametophyte and mean sporophyte TPM values ⍰ 1, log2(fold change) <1 or >-1 and/or adjusted *p*-value ⍰ 0.05) and unexpressed genes (mean gametophyte and mean sporophyte TPM values both <1).

### Gene ontology and prediction of subcellular localisation

Gene ontology term annotations were extracted from the Interproscan annotation files generated by the Phaeoexplorer consortium (https://phaeoexplorer.sb-roscoff.fr/downloads/; (Denoeud *et al*., 2024). The subcellular localisation of proteins was predicted by identifying amino-terminal targeting peptides using HECTAR version 1.3 (Gschloessl *et al*., 2008).

### Calculation of breadth of expression and Jaccard indices

Breadth of expression (tau) was calculated using a custom script in R based on the following formula:

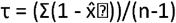

*Where* 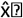 is the expression level of gene i normalized by the maximum expression value across all conditions (mean log_2_(NormalisedCounts +1)), and n is the number of conditions.

Jaccard indices were calculated and plotted using vegan version 2.6.8 (https://cran.r-project.org/web/packages/vegan/index.html).

### Enrichment analysis

Enrichment of co-expression modules in differentially-expressed genes, predicted subcellular localisations or gene ontology terms were computed using the enricher function from the R package clusterProfiler (Yu *et al*., 2012) version 4.2.2. *p*-values were adjusted for multiple testing based on the Benjamini-Hochberg false discovery rate.

### Identification of modules of co-expressed genes

For both the *Ectocarpus* species 7 and the *D. dichotoma* datasets, raw counts were normalised using the estimateSizeFactors and counts (with option normalized = TRUE) functions from the R package DESeq2 (Love *et al*., 2014) version 1.34.0. Genes with a normalised count below 45 in all the samples analysed for the relevant species were removed. The normalised counts matrix was then transformed with the log1p (=log_2_(x+1)) R function. To fit a scale-free topology model, the soft-thresholding power was selected using the pickSoftThreshold function, and gene modules were built using the blockwiseModules function from the WGCNA R package (Langfelder and Horvath, 2008) version 1.71. For *Ectocarpus* species 7, this function was run with options power = 16, maxBlockSize = 16100 (which yields a topological matrix in one block), minModuleSize = 30, reassignThreshold = 0, mergeCutHeight = 0.2, pamRespectsDendro = FALSE, randomSeed = T, corType = “Pearson”. For *D. dichotoma*, the function was run with power = 18, maxBlockSize = 16000 (which yields a topological matrix in one block), minModuleSize = 30, reassignThreshold = 0, mergeCutHeight = 0.25, pamRespectsDendro = FALSE, randomSeed = T, corType = “Pearson”. Complete module annotation and module membership (MM) scores (or kME, representing the correlation between the gene and the module eigengene) can be found, together with associated *p*-values, in Supplementary Tables S6 and S8. *D. dichotoma* gene module descriptions are in Figure S10B.

### Conservation of gene co-expression modules between *Ectocarpus* species 7 and *D. dichotoma*

An orthologue analysis was carried out specifically for *Ectocarpus* species 7 and *D. dichotoma* using Orthofinder (Emms and Kelly, 2019) version 2.5.2 to optimise the identification of one-to-one orthologues for these two species (Table S7). The one-to-one orthologue set generated by this analysis was then used to evaluate module conservation across the two species. The WGCNA package modulePreservation function (networkType = “signed”, nPermutations = 20, maxGoldModuleSize = 1000, maxModuleSize = 2500) and the NetRep package (Ritchie *et al*., 2016) version 1.2.7 modulePreservation function (nPerm=10000, on the WGCNA-computed correlation and TOM matrices) were used to compute module preservation statistics. In addition, the silhouette coefficient for modules was computed using the silhouette_SimilarityMatrix function from the

CancerSubtypes R package (Xu *et al*., 2017) version 1.20.0 on the correlation matrix. All values can be found in Table S8. The gene correlation heatmap (Figure 6B) was produced using the NetRep (Ritchie *et al*., 2016) version 1.2.7 plotCorrelation function. The gold module was a random sample of 1,000 *Ectocarpus* species 7 genes that provided a control distribution of gene expression patterns for comparisons with *D. dichotoma* modules. Cross-species module comparisons were visualized as a contingency table (Figure 6C) constructed by interpolating orthologous gene counts between the *Ectocarpus* species 7 and *D. dichotoma* datasets. For each pair of modules, a list of orthologues present in at least one of the two modules was established and a Fisher’s exact test was performed on Boolean vectors indicating whether each gene was present in each of the two modules to assess the statistical association between module assignment in both species. The objective was therefore to determine whether each pair of *Ectocarpus* species 7 and *D. dichotoma* modules shared a greater number of one-to-one orthologues than expected from a random distribution. Fisher’s exact test was applied using the fisher.test function from the R stats package (version 4.1.2). Cells in the contingency table were colour-coded according to -log_10_(*p*-value).

The contingency table (Figure 6C) was constructed by interpolating gene counts between the *Ectocarpus* species 7 and *D. dichotoma* datasets and using the fisher.test function (alternative = “greater”) from the R stats package (version 4.1.2).

### Detection of conservation of gene expression patterns between *Ectocarpus* species 7 and *D. dichotoma*

To assess gene expression pattern preservation between *Ectocarpus* species 7 and *D. dichotoma*, a z-score (mean for one condition – mean across all conditions) / standard deviation) was computed from the log_2_(NormalisedCounts+1) values for each of the six common timepoints between the *Ectocarpus* species 7 and *D. dichotoma* datasets (Table S1). A Pearson correlation coefficient was then calculated using the R stats package (version 4.1.2) cor.test function (alternative = “greater”, method = “pearson”).

### Manual reannotation of ribosomal protein genes and three-dimensional representation of ribosomal proteins

*Ectocarpus* species 7 ribosomal proteins were manually annotated using the recent *Arabidopsis thaliana* nomenclature (Scarpin *et al*., 2023) as a reference (Table S12). *Ectocarpus* species 7 ribosomal proteins were compared with the *A. thaliana* ribosomal protein dataset using BLASTp and the annotations transferred based on the best BLASTp match. To visualise differentially expression in relation to the structure of the ribosome, *Ectocarpus* species 7 ribosomal proteins were mapped onto the *Triticum aestivum* ribosome structure (Protein data bank model 4V7E; (Gogala *et al*., 2014) and the mapping was visualised using UCSF ChimeraX version 1.9 (Meng *et al*., 2023).

### General data treatment and figure preparation tools

Several R modules were used at multiple stages of the study. These included rtracklayer (Lawrence *et al*., 2009) version 1.52.1, Tidyverse (Wickham *et al*., 2019) version 2.0.0, and sjmisc (Lüdecke, 2018) version 2.8.9, which were used to handle and analyse data, and ggplot2 (Wickham *et al*., 2016) version 3.4.2 and Pheatmap (Kolde, 2019) version 1.0.12, which were used to generate figures.

## Data availability

All the new sequence data generated by this study have been deposited in the European Bioinformatics Institute/European Nucleotide Archive (EBI/ENA) database and are publicly available (see Table S1 for details).

### CRediT authorship contribution statement

Conceptualization: P.R., O.G., J.M.C.; Formal analysis: P.R., O.G.; Investigation: P.R., O.G.; Resources: B.N., J.-M.A; Data Curation: P.R., O.G.; Writing - Original Draft: P.R.; Writing - Review & Editing: all authors; Visualization: P.R., O.G.; Supervision: O.G., J.M.C.; Project administration: J.M.C.; Funding acquisition: : J.M.C.

## Acknowledgments

We would like to thank Julia Morales for helpful discussions about ribosomal proteins, Laurence Dartevelle and Elodie Rolland for help with algal cultures, Delphine Scornet, Masakazu Hoshino, Inka Bartsch, Akira F. Peters, Samuel Boscq, Shannon DeVanney for providing seaweed photographs. We are grateful to the Roscoff Bioinformatics platform ABiMS (http://abims.sb-roscoff.fr), which is part of the Institut Français de Bioinformatique (ANR-11-INBS-0013) and BioGenouest network, for providing both help and computing and storage resources.

## Supplementary Data

**Figure S1. Schematic life cycles and photographs of the sporophyte and gametophyte generations of brown algal species analysed in this study illustrating the different degrees of generational dimorphism**.

Size bars indicate the approximate sizes of each generation of each life cycle, providing an indication of the degree of dimorphism between the two generations. The life cycle and the morphology and size of the sporophyte and gametophyte generations of *Saccharina japonica* (not shown) are highly similar to those of *Saccharina latissima*. Photograph credits: *M. pyrifera* sporophyte, Shannon DeVanney; *S. latissima*, Samuel Boscq (Roscoff Biological Station); *M. clavaeformis*, Akira F. Peters (Bezhin Rosco); *S. promiscuus* sporophyte, Masakazu Hoshino (Kobe University); *P. littoralis* gametophyte, Inka Bartsch (Alfred Wegener Institut) and Delphine Scornet (Roscoff Biological Station); *P. littoralis* sporophyte, Alfred Wegener Institut and Delphine Scornet (Roscoff Biological Station); *S. polyschides* gametophyte, *M. pyrifera* gametophyte, *E. siliculosus*, Delphine Scornet (Roscoff Biological Station).

**Figure S2. Light microscopy images of *Ectocarpus* species 7 initial cell and elongating initial cell**

A. *Ectocarpus* species 7 strain Ec32 initial cell (settled male Ec32 gamete, 24 h after release). B. *Ectocarpus* species 7 strain Ec32 elongating initial cell (settled male gamete, 48 h after release). Inverted Leica microscope, magnification: x200, scale bar: 100 µm.

**Figure S3. Breadth of expression (tau) of genes that were differentially expressed, or not, across the transition from gamete to sporophyte initial cell**.

Significantly downregulated and upregulated genes were identified using DESeq2. Dotted lines indicate mean tau values.

**Figure S4. Hierarchical cluster trees showing co-expression modules identified using WGCNA**.

**A**. *Ectocarpus* species 7. **B**. *D. dichotoma*. Grouped branches correspond to modules, which are indicated by coloured boxes underneath the tree.

**Figure S5. Enriched GO terms in *Ectocarpus* species 7 gene modules**

Statistically-significant enriched GO terms in *Ectocarpus* species 7 gene modules. Enrichment is indicated as log2 of the ratio of the proportion of genes assigned to the GO term in the module divided by the proportion for the whole genome. When it was possible to manually assign a general function to a module, the annotation is indicated in brackets after the module name. padj, *p*-value adjusted for multiple testing based on the Benjamini-Hochberg false discovery rate.

**Figure S6. Transcript abundance heatmap for *Ectocarpus* species 7 CAZYme genes**

Heatmaps with log_2_(NormalisedCounts+1) for replicate samples of (from left to right) free-swimming male gamete, sporophyte initial cell (24 hours after gamete release), elongating sporophyte initial cell (48 hours after gamete release), sporophyte 2-5 cell stage, non-fertile adult sporophyte, fertile sporophyte, non-fertile female and male gametophyte, fertile female and male gametophyte *Ectocarpus* species 7 developmental stages for GT23 and PL41 annotated genes. Left annotation track: differential expression (DEG) analysis results in the transitions between (1) free-swimming male gamete and sporophyte initial cell stage, and (2) adult sporophyte and gametophyte; right annotation track: WGCNA module colours.

**Figure S7. Transcript abundance heatmap for *Ectocarpus* species 7 transcription factor genes that have one-to-one *D. dichotoma* orthologues**

Hierarchically clustered heatmap of log_2_(NormalisedCounts+1) for 325 *Ectocarpus* species 7 TAP genes (Denoeud *et al*., 2024) and 90 EsV1-7 domain genes (Macaisne *et al*., 2017) for replicate samples of the following life cycle stages: (from left to right) free-swimming male gamete, sporophyte initial cell (24 hours after gamete release), elongating sporophyte initial cell (48 hours after gamete release), sporophyte 2-5 cell stage, non-fertile adult sporophyte, fertile sporophyte, non-fertile female and male gametophytes, fertile female and male gametophytes. Left annotation track: differential expression (DEG) analysis results in the transitions between 1) free-swimming male gamete and initial cell stage, and 2) sporophyte and gametophyte; right annotation track: WGCNA module colours.

**Figure S8. Transcript abundance heatmap for *Ectocarpus* species 7 ribosomal genes**

Heatmap with log_2_(NormalisedCounts+1) for (from left to right) free-swimming male gamete, sporophyte initial cell (24 hours after gamete release), elongating sporophyte initial cell (48 hours after gamete release), sporophyte 2-5 cell stage, non-fertile adult sporophyte, fertile sporophyte, non-fertile female and male gametophyte, fertile female and male gametophyte *Ectocarpus* species 7 developmental stages for ribosomal protein annotated genes. Left annotation track: differential expression analysis results in the transitions between (1) free-swimming male gamete and sporophyte initial cell stage, and (2) adult sporophyte and gametophyte; right annotation track: WGCNA module colours.

**Figure S9. Three dimensional representations of a ribosome illustrating differential expression of ribosomal protein genes**

A., B. Lateral views. C. Top view. D. Bottom view. E. and F. Intersubunit views of the small and large subunit, respectively. Small and large subunit ribosomal proteins are coloured in red and purple, respectively. Darker colour indicates that the gene encoding a protein was significantly upregulated between the gamete and sporophyte initial cell stages. Proteins coloured in white are present in the *Triticum aestivum* ribosome, which was used as a reference, but no equivalent was found in *Ectocarpus* species 7. RNAs are shown as ribbons, ribosomal RNAs in grey, messenger and transfer RNAs in green. A, P and T indicate the aminoacyl, peptidyl and exit tRNA binding sites, respectively. En, mRNA entry site; Ex, mRNA exit site; Tu, nascent peptide tunnel exit.

**Figure S10. *D. dichotoma* PCA and modules**

**A**. Principal component analysis of *D. dichotoma* gene expression across seven life cycle stages. **B**. Characterisation of *D. dichotoma* gene co-expression modules generated using WGCNA. Left to right: manually-assigned functional categories based on GO term enrichment; number of genes in the module positively or negatively correlated with the module eigengene; average module gene expression profiles for WGCNA modules across sperm (1 h after release), egg cell (15 min after release), zygote (1h after release), embryo (8h after fertilisation), fertile sporophyte, fertile female and male gametophyte life cycle stages; module eigengene dendrogram. The average module gene expression profiles were calculated using genes with a WGCNA MM >0.86.

**Figure S11. Enriched GO terms *D. dichotoma* gene modules**

Statistically-significant enriched GO terms in *D. dichotoma* gene modules. Enrichment is indicated as log2 of the ratio of the proportion of genes assigned to the GO term in the module divided by the proportion for the whole genome. padj, *p*-value adjusted for multiple testing based on the Benjamini-Hochberg false discovery rate.

**Figure S12. Transcript abundance heatmap for one-to-one orthologous transcription factors between *Ectocarpus* species 7 and *D. dichotoma***

Heatmap representing expression of one-to-one orthologous genes from *Ectocarpus* species 7 and *D. dichotoma* presented as the z-score of the log_2_(NormalisedCounts+1) for the following equivalent developmental stages in *Ectocarpus* species 7 and *D. dichotoma*: sperm, zygote, early sporophyte, adult sporophyte, female and male gametophyte. The lefthand tracks show Pearson correlation coefficients and log_10_ Benjamini-Hochberg-adjusted *p*-values (log10 padj).

**Table S1. RNA-seq data used in this study**

GA, gametophyte; SP, sporophyte; pSP, partheno-sporophyte; GBG, generation-biased genes.

**Table S2. Generation-biased and generation-specific gene sets in ten species of brown algae based on comparisons of adult sporophyte and gametophyte generations**.

**A**. Overview of the generation-biased and generation-specific gene sets of each species. **B**. Complete proteomes of the ten species indicating the results of the comparison of adult sporophytes and gametophytes, including log_2_(fold change), *p*-value, mean TPM and assigned generation-biased expression class. TPM, transcripts per kilobase million.

**Table S3. List of orthogroups for the ten brown algal species analysed for generation-biased gene expression**.

The genomes analysed (with ENA accession numbers or links to ORCAE in brackets) correspond to: *Dictyota dichotoma* strain ODC1387m (GCA_964200555), *Ectocarpus* species 7 strain Ec32 (https://bioinformatics.psb.ugent.be/orcae/overview/EctsiV2), *Macrocystis pyrifera* strain P11B4 (GCA_964200385), *Myriotrichia clavaeformis* strain Myr cla04 (GCA_964200105), *Pylaiella littoralis* strain U1.48 (GCA_964200295), *Saccharina latissima* strain SLPER63f7 (GCA_964200175), *Saccharina japonica* strain Ja (https://bioinformatics.psb.ugent.be/orcae/overview/Sacja), *Saccorhiza polyschides* strain SpolBR94m (GCA_964200605), *Scytosiphon promiscuus* strain Ot110409-Otamoi-16-male (GCA_964200365) and *Sphacelaria rigidula* strain Sph rig Cal Mo 4-1-68b (GCA_964200075). The one-to-one orthologues correspond to the 9,317 orthogroups identified using the relaxed criteria described in the Material and Methods.

**Table S4. Counts of RNA-seq reads mapped to *Ectocarpus* species 7 genes**

This data was used to calculate the gene expression levels reported in Table S4.

**Table S5. Functional and expression-related information for *Ectocarpus* species 7 genes**

List of all *Ectocarpus* species 7 genes with transcript abundance under each of the conditions analysed, measured as log_2_(NormalisedCounts +1), WGCNA module assignment (Colour), WGCNA module membership (MM) scores and associated *p*-values (pMM) for each gene in each module, the functional annotation for each gene (description), the manually assigned functional category (Manual.functional.category), the DESeq2 output for the comparison between free-swimming male gamete and sporophyte initial cell stage (gamete_initial), and adult sporophyte and gametophyte (sporophyte_gametophyte) with the corresponding differential expression (DE) annotation, the HECTAR output for each gene with the predicted targeting category and corresponding scores.

Table S6. Overview of the gene co-expression modules for *Ectocarpus* species 7 and *D. dichotoma* The grey modules contain all the genes that could not be assigned to a co-expression module. The gold module was a random sample of 1,000 *Ectocarpus* species 7 genes that provided a null distribution of gene expression patterns for the comparisons with *D. dichotoma* modules.

**Table S7. Orthofinder orthogroup analysis of *Ectocarpus* species 7 and *D. dichotoma***.

The genomes analysed (with ENA accession numbers or links to ORCAE in brackets) correspond to: *Dictyota dichotoma* strain ODC1387m (GCA_964200555) and *Ectocarpus* species 7 strain Ec32 (https://bioinformatics.psb.ugent.be/orcae/overview/EctsiV2).

**Table S8. Conservation of life-cycle-related co-expression modules between *Ectocarpus* species 7 and *D. dichotoma***

For each *Ectocarpus* species 7 module: preservation statistics computed by the WGCNA modulePreservation function (200 permutations), preservation statistics computed by the NetRep modulePreservation function (10,000 permutations) with the associated log_10_(*p*-value) (Bonferroni-corrected for WGCNA), and the silhouette score for each module in *D. dichotoma*. WGCNA - medianRank statistic, aggregation of the densityMedianRank and connectivityMedianRank statistics; WGCNA – medianRankDensity.pres, WGCNA density median rank preservation statistic; WGCNA – medianRankConnectivity.pres, WGCNA connectivity median rank preservation statistic; WGCNA - propVarExplained.pres, coherence: proportion of module variance explained by the module eigengene (=summary profile = eigenvector of the 1st principal component across all observations for every node composing the module); WGCNA – meanSignAwareKME.pres, average node contribution: average Pearson correlation coefficient to the module’s summary profile; WGCNA - meanSignAwareCorDat.pres, density of correlation structure: how strongly modules are correlated in the test dataset, average node correlation; WGCNA - meanAdj.pres, average edge weight: average connection strength between nodes; WGCNA – meanMAR.pres, average module adjacency ratio; WGCNA: cor.kIM, concordance of intramodular connectivity: similarity of the relative rank of each nodes’ weighted degree (intramodular connectivity) across datasets; WGCNA – cor.kME, concordance of node contribution: preservation of relative rank of nodes (ordered by Pearson correlation coefficient to the module’s summary profile) across datasets; WGCNA – cor.kMEall; concordance of node contribution: preservation of relative rank of nodes (ordered by Pearson correlation coefficient to the module’s summary profile) across datasets taking into account the kME of the gene for all modules; WGCNA - cor.cor, concordance of correlation structure: quantifies how similar the correlation structure is across datasets; WGCNA – cor.MAR, concordance of module adjacency ratio: concordance of the ration between the maximum and the minimum adjacency in the module; WGCNA - separability.pres, separability preservation; NetRep - avg.weight, average edge weight: average connection strength between nodes; NetRep - coherence, coherence, i. e. proportion of module variance explained by the module eigengene (=summary profile = eigenvector of the 1st principal component across all observations for every node composing the module); NetRep – cor.cor, concordance of correlation structure: quantifies how similar the correlation structure is across datasets; NetRep - cor.degree, concordance of intramodular connectivity: similarity of the relative rank of each nodes’ weighted degree (intramodular connectivity) across datasets; NetRep - cor.contrib, concordance of node contribution: preservation of relative rank of nodes (ordered by Pearson correlation coefficient to the module’s summary profile) across datasets; NetRep - avg.cor, density of correlation structure: how strongly modules are correlated in the test dataset, average node correlation; NetRep - avg.contrib, average node contribution: average Pearson correlation coefficient to the module’s summary profile; silhouette.cluster.width, Silhouette coefficient; silhouette.distance.to.average, Silhouette coefficient. The “gold” module was a random sample of 1,000 *Ectocarpus* species 7 genes that provided a control distribution of gene expression patterns for comparisons with *D. dichotoma* modules. The “darkturquoise” module was not included in this analysis because it contained zero orthologues (Table S5).

**Table S9. Counts of RNA-seq reads mapped to *D. dichotoma* genes**

This data was used to calculate the gene expression levels reported in Table S8.

**Table S10. Functional and expression-related information for *D. dichotoma* genes**

List of all *D. dichotoma* genes with the log_2_(NormalisedCounts+1) under each condition used for the analysis, WGCNA module assignment (Colour), WGCNA module membership (MM) scores and associated *p*-values (pMM) for each gene in each module.

**Table S11. Correlation between the expression patterns of orthologous transcription factors in *Ectocarpus* species 7 and *D. dichotoma***

Correlation was evaluated based on comparisons six approximately equivalent stages for the two species (Table S1). The names of the *Ectocarpus* species 7 and *D. dichotoma* genes in each orthogroup are given in Figure S12. padj, *p*-value adjusted for multiple testing based on the Benjamini-Hochberg false discovery rate.

**Table S12. Manual re-annotation of the *Ectocarpus* species 7 ribosomal proteins**.

The e-values and scores are for BLASTp queries of the *Ectocarpus* species 7 ribosomal proteins against the *Arabidopsis thaliana* ribosomal protein dataset.

## References

Ahmed S, Cock JM, Pessia E, Luthringer R, Cormier A, Robuchon M, Sterck L, Peters AF, Dittami SM, Corre E, Valero M, Aury JM, Roze D, Van de Peer Y, Bothwell J, Marais GA, Coelho SM (2014) A Haploid System of Sex Determination in the Brown Alga Ectocarpus sp. Curr Biol 24:1945– 1957.

Akita S, Vieira C, Hanyuda T, Rousseau F, Cruaud C, Couloux A, Heesch S, Cock JM, Kawai H (2022) Providing a phylogenetic framework for trait-based analyses in brown algae: Phylogenomic tree inferred from 32 nuclear protein-coding sequences. Molecular Phylogenetics and Evolution 168:107408.

Al-Hinai TZS, Mackay CL, Fry SC (2024) Fruit softening: evidence for rhamnogalacturonan lyase action in vivo in ripe fruit cell walls. Annals of Botany 133:547–558.

Anderson SN, Johnson CS, Chesnut J, Jones DS, Khanday I, Woodhouse M, Li C, Conrad LJ, Russell SD, Sundaresan V (2017) The Zygotic Transition Is Initiated in Unicellular Plant Zygotes with Asymmetric Activation of Parental Genomes. Dev Cell 43:349-358.e4.

Andrews S (2016) FastQC A Quality Control tool for High Throughput Sequence Data. http://www.bioinformatics.babraham.ac.uk/projects/fastqc/.

Arun A, Coelho SM, Peters AF, Bourdareau S, Pérès L, Scornet D, Strittmatter M, Lipinska AP, Yao H, Godfroy O, Montecinos GJ, Avia K, Macaisne N, Troadec C, Bendahmane A, Cock JM (2019) Convergent recruitment of TALE homeodomain life cycle regulators to direct sporophyte development in land plants and brown algae. Elife 8:e43101.

Arun A, Peters NT, Scornet D, Peters AF, Cock JM, Coelho SM (2013) Non-cell autonomous regulation of life cycle transitions in the model brown alga Ectocarpus. New Phytol 197:503–510.

Bacher R, Leng N, Chu L-F, Ni Z, Thomson JA, Kendziorski C, Stewart R (2018) Trendy: segmented regression analysis of expression dynamics in high-throughput ordered profiling experiments. BMC Bioinformatics 19:380.

Bogaert KA, Beeckman T, De Clerck O (2017) Two-step cell polarization in algal zygotes. Nat Plants 3:16221.

Bourdareau S, Tirichine L, Lombard B, Loew D, Scornet D, Wu Y, Coelho SM, Cock JM (2021) Histone modifications during the life cycle of the brown alga Ectocarpus. Genome Biology 22:12.

Bringloe TT, Starko S, Wade RM, Vieira C, Kawai H, Clerck OD, Cock JM, Coelho SM, Destombe C, Valero M, Neiva J, Pearson GA, Faugeron S, Serrão EA, Verbruggen H (2020) Phylogeny and Evolution of the Brown Algae. Critical Reviews in Plant Sciences 39:281–321.

Carney LT, Edwards MS (2006) Cryptic Processes in the Sea: A Review of Delayed Development in the Microscopic Life Stages of Marine Macroalgae. Algae 21:161–168.

Carvalho-Santos Z, Azimzadeh J, Pereira-Leal JB, Bettencourt-Dias M (2011) Evolution: Tracing the origins of centrioles, cilia, and flagella. J Cell Biol 194:165–175.

Chen J, Strieder N, Krohn NG, Cyprys P, Sprunck S, Engelmann JC, Dresselhaus T (2017) Zygotic Genome Activation Occurs Shortly after Fertilization in Maize. The Plant Cell 29:2106–2125.

Choi S-W, Graf L, Choi JW, Jo J, Boo GH, Kawai H, Choi CG, Xiao S, Knoll AH, Andersen RA, Yoon HS (2024) Ordovician origin and subsequent diversification of the brown algae. Curr Biol:S0960-9822(23)01769-4.

Choksi SP, Lauter G, Swoboda P, Roy S (2014) Switching on cilia: transcriptional networks regulating ciliogenesis. Development 141:1427–1441.

Chu JSC, Baillie DL, Chen N (2010) Convergent evolution of RFX transcription factors and ciliary genes predated the origin of metazoans. BMC Evol Biol 10:130.

Cock JM et al. (2010) The Ectocarpus genome and the independent evolution of multicellularity in brown algae. Nature 465:617–621.

Cock JM, Godfroy O, Macaisne N, Peters AF, Coelho SM (2014) Evolution and regulation of complex life cycles: a brown algal perspective. Curr Opin Plant Biol 17:1–6.

Coelho SM, Cock JM (2020) Brown algal model organisms. Ann Rev Genet 54:71–92.

Coelho SM, Godfroy O, Arun A, Le Corguillé G, Peters AF, Cock JM (2011) OUROBOROS is a master regulator of the gametophyte to sporophyte life cycle transition in the brown alga Ectocarpus. Proc Natl Acad Sci USA 108:11518–11523.

Coelho SM, Scornet D, Rousvoal S, Peters N, Dartevelle L, Peters AF, Cock JM (2012a) Genetic crosses between Ectocarpus strains. Cold Spring Harb Protoc 2012:262–265.

Coelho SM, Scornet D, Rousvoal S, Peters NT, Dartevelle L, Peters AF, Cock JM (2012b) How to cultivate Ectocarpus. Cold Spring Harb Protoc 2012:258–261.

Cormier A, Avia K, Sterck L, Derrien T, Wucher V, Andres G, Monsoor M, Godfroy O, Lipinska A, Perrineau M-M, Van De Peer Y, Hitte C, Corre E, Coelho SM, Cock JM (2017) Re-annotation, improved large-scale assembly and establishment of a catalogue of noncoding loci for the genome of the model brown alga Ectocarpus. New Phytol 214:219–232.

Couceiro L, Le Gac M, Hunsperger HM, Mauger S, Destombe C, Cock JM, Ahmed S, Coelho SM, Valero M, Peters AF (2015) Evolution and maintenance of haploid-diploid life cycles in natural populations: The case of the marine brown alga Ectocarpus. Evolution 69:1808–1822.

Danecek P, Bonfield JK, Liddle J, Marshall J, Ohan V, Pollard MO, Whitwham A, Keane T, McCarthy SA, Davies RM, Li H (2021) Twelve years of SAMtools and BCFtools. Gigascience 10:giab008.

Denoeud F et al. (2024) Evolutionary genomics of the emergence of brown algae as key components of coastal ecosystems. Cell 187:6943–6965.

Dopler A et al. (2024) P-stalk ribosomes act as master regulators of cytokine-mediated processes. Cell Available at: https://www.sciencedirect.com/science/article/pii/S0092867424011395 [Accessed November 25, 2024].

Eger AM, Marzinelli EM, Beas-Luna R, Blain CO, Blamey LK, Byrnes JEK, Carnell PE, Choi CG, Hessing-Lewis M, Kim KY, Kumagai NH, Lorda J, Moore P, Nakamura Y, Pérez-Matus A, Pontier O, Smale D, Steinberg PD, Vergés A (2023) The value of ecosystem services in global marine kelp forests. Nat Commun 14:1894.

Emms DM, Kelly S (2019) OrthoFinder: phylogenetic orthology inference for comparative genomics. Genome Biol 20:238.

Godfroy O, Uji T, Nagasato C, Lipinska AP, Scornet D, Peters AF, Avia K, Colin S, Mignerot L, Motomura T, Cock JM, Coelho SM (2017) DISTAG/TBCCd1 Is Required for Basal Cell Fate Determination in Ectocarpus. Plant Cell 29:3102–3122.

Godfroy O, Zheng M, Yao H, Henschen A, Peters AF, Scornet D, Colin S, Ronchi P, Hipp K, Nagasato C, Motomura T, Cock JM, Coelho SM (2023) The baseless mutant links protein phosphatase 2A with basal cell identity in the brown alga Ectocarpus. Development 150:dev201283.

Gogala M, Becker T, Beatrix B, Armache J-P, Barrio-Garcia C, Berninghausen O, Beckmann R (2014) Structures of the Sec61 complex engaged in nascent peptide translocation or membrane insertion. Nature 506:107–110.

Graf L, Shin Y, Yang JH, Hwang IK, Yoon HS (2022) Transcriptome analysis reveals the spatial and temporal differentiation of gene expression in the sporophyte of Undaria pinnatifida. Algal Research 68:102883.

Gschloessl B, Guermeur Y, Cock J (2008) HECTAR: a method to predict subcellular targeting in heterokonts. BMC Bioinf 9:393.

Inoue A, Ojima T (2019) Functional identification of alginate lyase from the brown alga Saccharina japonica. Sci Rep 9:4937.

Kim D, Paggi JM, Park C, Bennett C, Salzberg SL (2019) Graph-based genome alignment and genotyping with HISAT2 and HISAT-genotype. Nat Biotechnol 37:907–915.

Kolde R (2019) Pheatmap: pretty heatmaps. R package version 1:726.

Krueger F (2015) Trim Galore!: A wrapper around Cutadapt and FastQC to consistently apply adapter and quality trimming to FastQ files, with extra functionality for RRBS data. Babraham Institute Available at: https://cir.nii.ac.jp/crid/1370294643762929691 [Accessed October 8, 2024].

Langfelder P, Horvath S (2008) WGCNA: an R package for weighted correlation network analysis. BMC Bioinformatics 9:559.

Lawrence M, Gentleman R, Carey V (2009) rtracklayer: an R package for interfacing with genome browsers. Bioinformatics 25:1841–1842.

Li L, Tian G, Peng H, Meng D, Wang L, Hu X, Tian C, He M, Zhou J, Chen L, Fu C, Zhang W, Hu Z (2018) New class of transcription factors controls flagellar assembly by recruiting RNA polymerase II in Chlamydomonas. Proc Natl Acad Sci U S A 115:4435–4440.

Liang Z, Wang X, Zhang P, Liu W, Wang W, Liu F (2023) Chronological development of the morphological, physiological, biochemical, and transcriptomic changes provides insights into the mechanisms of gametogenesis in Saccharina japonica. J Appl Phycol 35:785–802.

Liao Y, Smyth GK, Shi W (2014) featureCounts: an efficient general purpose program for assigning sequence reads to genomic features. Bioinformatics 30:923–930.

Lipinska AP, Serrano-Serrano ML, Cormier A, Peters AF, Kogame K, Cock JM, Coelho SM (2019) Rapid turnover of life-cycle-related genes in the brown algae. Genome Biol 20:35.

Love MI, Huber W, Anders S (2014) Moderated estimation of fold change and dispersion for RNA-seq data with DESeq2. Genome Biol 15:550.

Lüdecke D (2018) sjmisc: Data and Variable Transformation Functions. Journal of Open Source Software 3:754.

Macaisne N, Liu F, Scornet D, Peters AF, Lipinska A, Perrineau M-M, Henry A, Strittmatter M, Coelho SM, Cock JM (2017) The Ectocarpus IMMEDIATE UPRIGHT gene encodes a member of a novel family of cysteine-rich proteins with an unusual distribution across the eukaryotes. Development 144:409–418.

Martin DE, Soulard A, Hall MN (2004) TOR regulates ribosomal protein gene expression via PKA and the Forkhead transcription factor FHL1. Cell 119:969–979.

Mazéas L, Bouguerba-Collin A, Cock JM, Denoeud F, Godfroy O, Brillet-Guéguen L, Barbeyron T, Lipinska AP, Delage L, Corre E, Drula E, Henrissat B, Czjzek M, Terrapon N, Hervé C (2024) Candidate genes involved in biosynthesis and degradation of the main extracellular matrix polysaccharides of brown algae and their probable evolutionary history. BMC Genomics 25:950.

Méndez-Yañez A, González M, Carrasco-Orellana C, Herrera R, Moya-León MA (2020) Isolation of a rhamnogalacturonan lyase expressed during ripening of the Chilean strawberry fruit and its biochemical characterization. Plant Physiology and Biochemistry 146:411–419.

Meng EC, Goddard TD, Pettersen EF, Couch GS, Pearson ZJ, Morris JH, Ferrin TE (2023) UCSF ChimeraX: Tools for structure building and analysis. Protein Science 32:e4792.

Müller DG (1967) Generationswechsel, Kernphasenwechsel und Sexualität der Braunalge Ectocarpus siliculosus im Kulturversuch. Planta 75:39–54.

Ni C, Buszczak M (2023) The homeostatic regulation of ribosome biogenesis. Semin Cell Dev Biol 136:13–26.

Pearson GA, Martins N, Madeira P, Serrão EA, Bartsch I (2019) Sex-dependent and -independent transcriptional changes during haploid phase gametogenesis in the sugar kelp Saccharina latissima. PLOS ONE 14:e0219723.

Peters AF, Scornet D, Ratin M, Charrier B, Monnier A, Merrien Y, Corre E, Coelho SM, Cock JM (2008) Life-cycle-generation-specific developmental processes are modified in the immediate upright mutant of the brown alga Ectocarpus siliculosus. Development 135:1503–1512.

Ritchie SC, Watts S, Fearnley LG, Holt KE, Abraham G, Inouye M (2016) A Scalable Permutation Approach Reveals Replication and Preservation Patterns of Network Modules in Large Datasets. cels 3:71–82.

Scarpin MR, Busche M, Martinez RE, Harper LC, Reiser L, Szakonyi D, Merchante C, Lan T, Xiong W, Mo B, Tang G, Chen X, Bailey-Serres J, Browning KS, Brunkard JO (2023) An updated nomenclature for plant ribosomal protein genes. The Plant Cell 35:640–643.

Shan T, Li Q, Wang X, Pang S (2020) Full-length transcriptome sequencing and comparative transcriptomic analysis of different developmental stages of the sporophyll in Undaria pinnatifida (Laminariales: Alariaceae). J Appl Phycol 32:2081–2092.

Shao Z, Zhang P, Lu C, Li S, Chen Z, Wang X, Duan D (2019) Transcriptome sequencing of Saccharina japonica sporophytes during whole developmental periods reveals regulatory networks underlying alginate and mannitol biosynthesis. BMC Genomics 20:975.

Starr RC, Zeikus JA (1993) UTEX-The culture collection of algae at the University of Texas at Austin 1993 list of cultures. J Phycol 29 (Suppl.):1–106.

Sterck L, Billiau K, Abeel T, Rouzé P, Van de Peer Y (2012) ORCAE: online resource for community annotation of eukaryotes. Nat Methods 9:1041.

Tadros W, Lipshitz HD (2009) The maternal-to-zygotic transition: a play in two acts. Development 136:3033–3042.

Tronholm A, Sansón M, Afonso-Carrillo J, De Clerck O (2008) Distinctive morphological features, life-cycle phases and seasonal variations in subtropical populations of Dictyota dichotoma (Dictyotales, Phaeophyceae). Bot Mar 51:132–144.

Uji T, Kandori T, Mizuta H (2024) Identification of differential gene expression related to reproduction in the sporophytes of Saccharina japonica. Front Plant Sci 15 Available at: https://www.frontiersin.org/journals/plant-science/articles/10.3389/fpls.2024.1417582/full [Accessed November 20, 2024].

Uluisik S, Seymour GB (2020) Pectate lyases: Their role in plants and importance in fruit ripening. Food Chemistry 309:125559.

van den Hoek C, Mann DG, Jahns HM (1995) Algae: An Introduction to Phycology. Cambridge: Cambridge University Press.

Vastenhouw NL, Cao WX, Lipshitz HD (2019) The maternal-to-zygotic transition revisited. Development 146:dev161471.

Wang P, Hendron R-W, Kelly S (2017) Transcriptional control of photosynthetic capacity: conservation and divergence from Arabidopsis to rice. New Phytol 216:32–45.

Waters MT, Wang P, Korkaric M, Capper RG, Saunders NJ, Langdale JA (2009) GLK transcription factors coordinate expression of the photosynthetic apparatus in Arabidopsis. Plant Cell 21:1109–1128.

Wawiórka L, Molestak E, Szajwaj M, Michalec-Wawiórka B, Mołoń M, Borkiewicz L, Grela P, Boguszewska A, Tchórzewski M (2017) Multiplication of Ribosomal P-Stalk Proteins Contributes to the Fidelity of Translation. Molecular and Cellular Biology 37:e00060–17.

Wickham H, Averick M, Bryan J, Chang W, McGowan LD, François R, Grolemund G, Hayes A, Henry L, Hester J (2019) Welcome to the Tidyverse. Journal of open source software 4:1686.

Wickham H, Chang W, Wickham MH (2016) Package ‘ggplot2.’ Create elegant data visualisations using the grammar of graphics Version 2:1–189.

Xu T, Le TD, Liu L, Su N, Wang R, Sun B, Colaprico A, Bontempi G, Li J (2017) CancerSubtypes: an R/Bioconductor package for molecular cancer subtype identification, validation and visualization. Bioinformatics 33:3131–3133.

Xue Z, Huang K, Cai C, Cai L, Jiang C, Feng Y, Liu Z, Zeng Q, Cheng L, Sun YE, Liu J, Horvath S, Fan G (2013) Genetic programs in human and mouse early embryos revealed by single-cell RNA sequencing. Nature 500:593–597.

Yao H, Scornet D, Jam M, Hervé C, Potin P, Correia LO, Coelho SM, Cock JM (2021) Biochemical characteristics of a diffusible factor that induces gametophyte to sporophyte switching in the brown alga Ectocarpus. Journal of Phycology 57:742–753.

Yu G, Wang L-G, Han Y, He Q-Y (2012) clusterProfiler: an R package for comparing biological themes among gene clusters. OMICS 16:284–287.

Zhang J, Li Y, Luo S, Cao M, Zhang L, Li X (2021) Differential gene expression patterns during gametophyte development provide insights into sex differentiation in the dioicous kelp Saccharina japonica. BMC Plant Biol 21:335.

Zhao P, Zhou X, Shen K, Liu Z, Cheng T, Liu D, Cheng Y, Peng X, Sun M-X (2019) Two-Step Maternal-to-Zygotic Transition with Two-Phase Parental Genome Contributions. Dev Cell 49:882-893.e5.

