## Supplementary figures and images for "Life-cycle-related gene expression patterns in the brown algae"

### Fig. S1

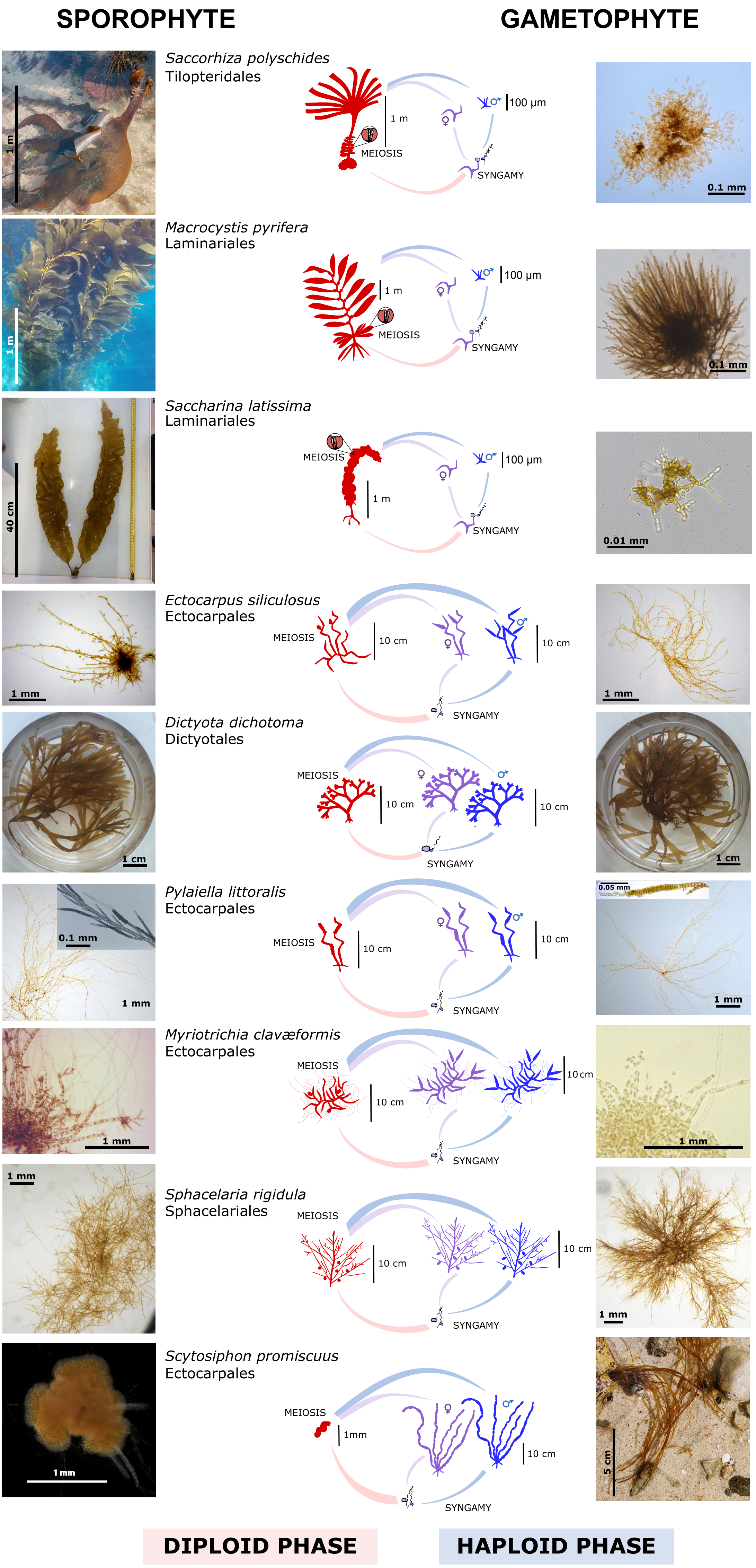

### Fig. S2

**A**

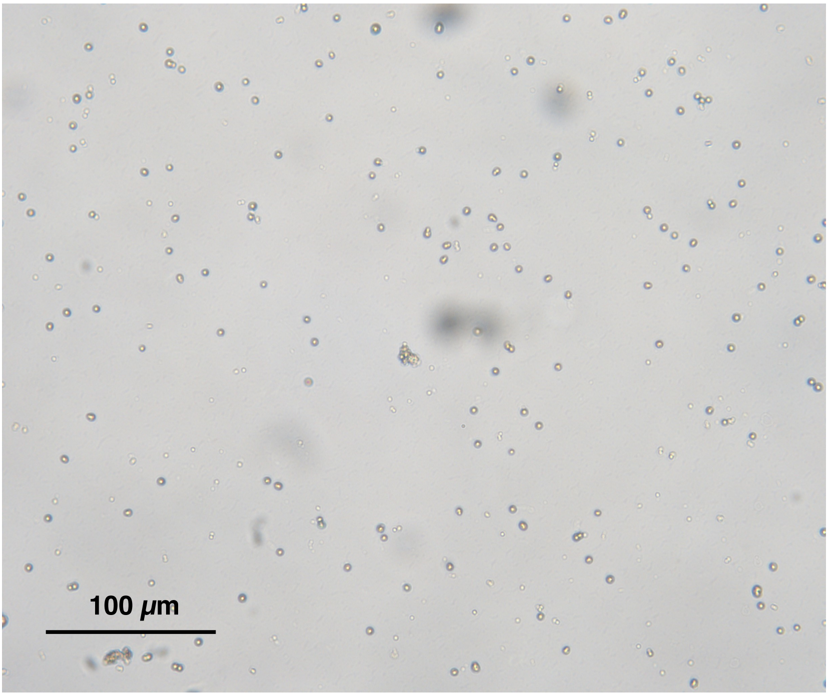


**B**

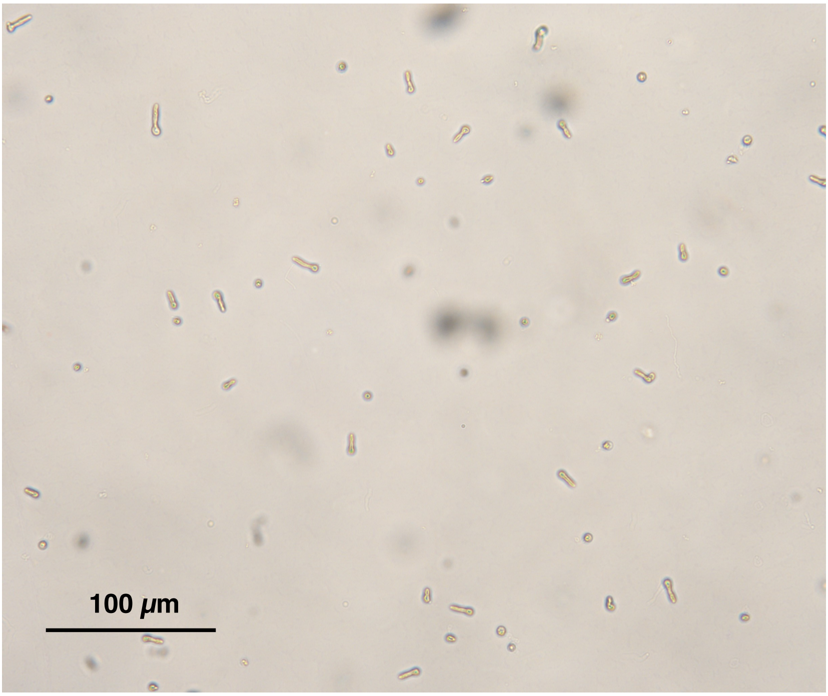

### Fig. S3

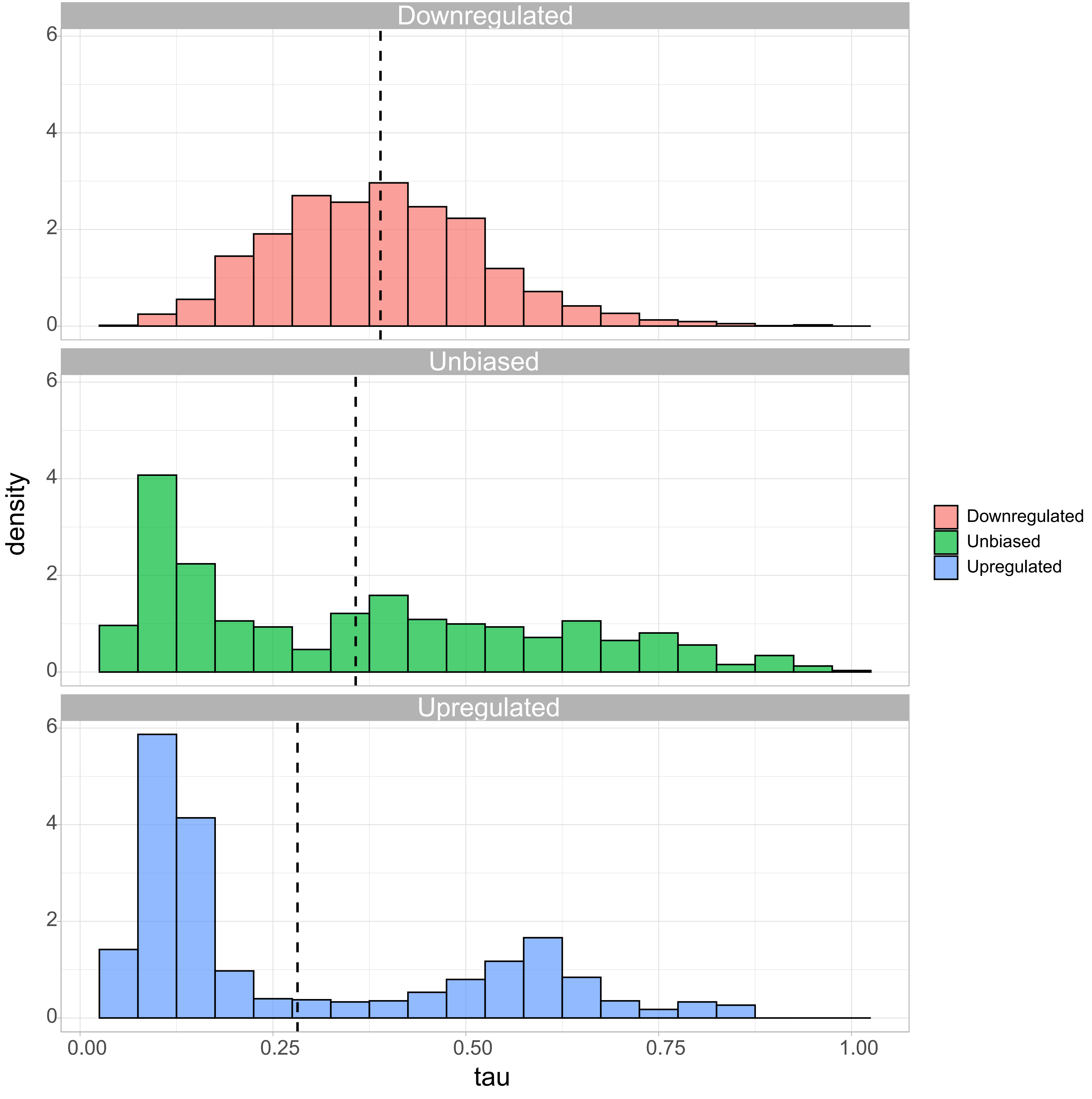

### Fig. S4

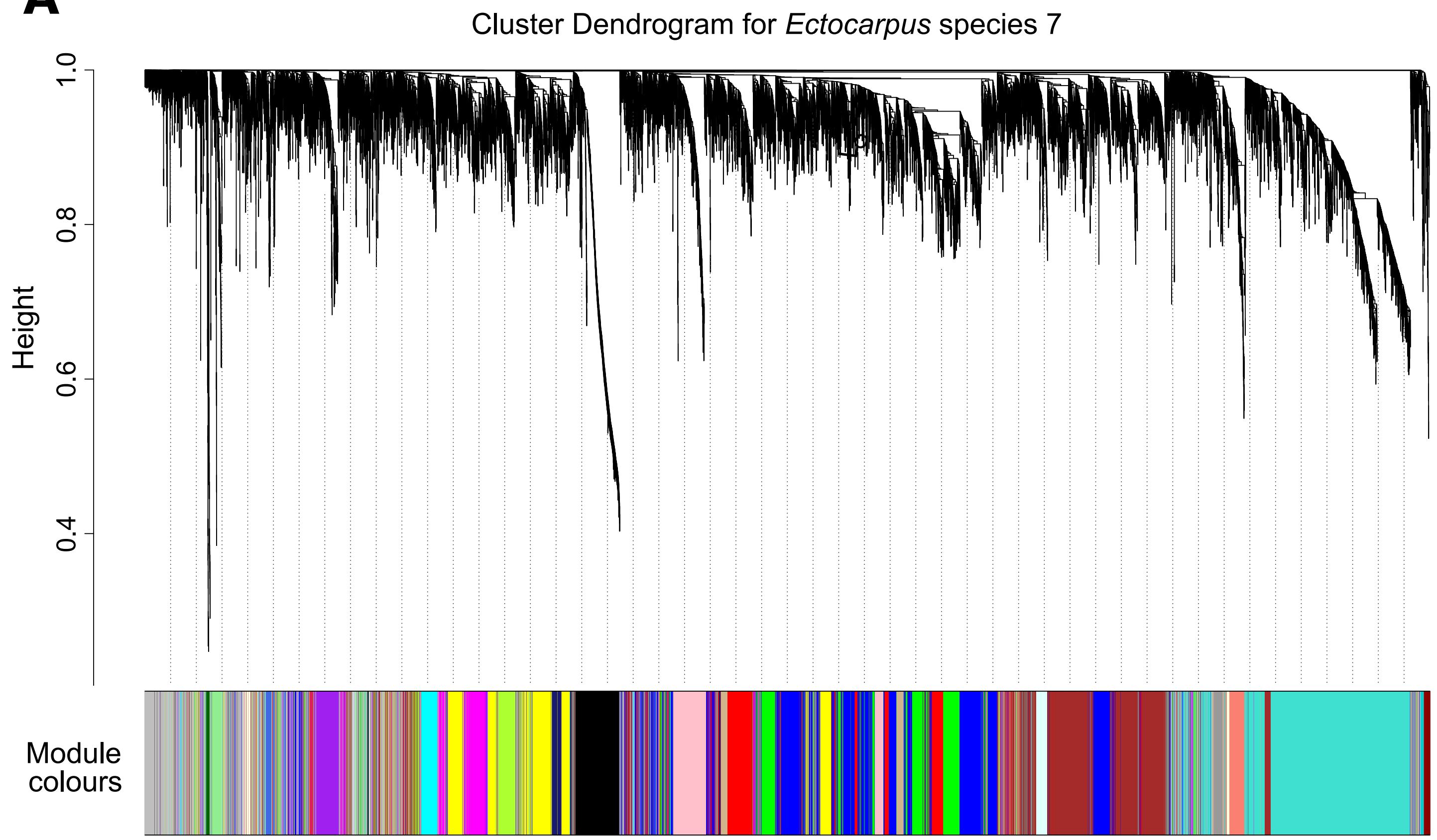

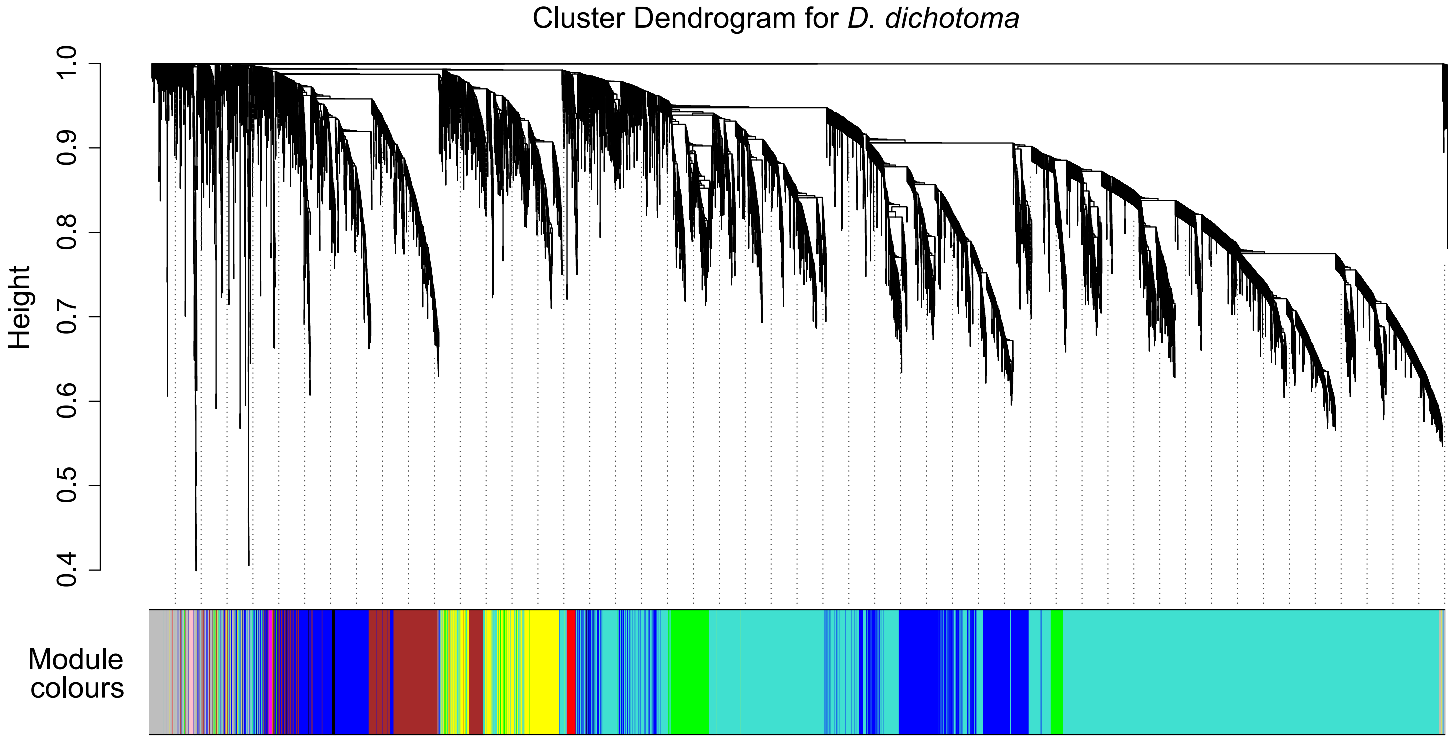

### Fig. S5

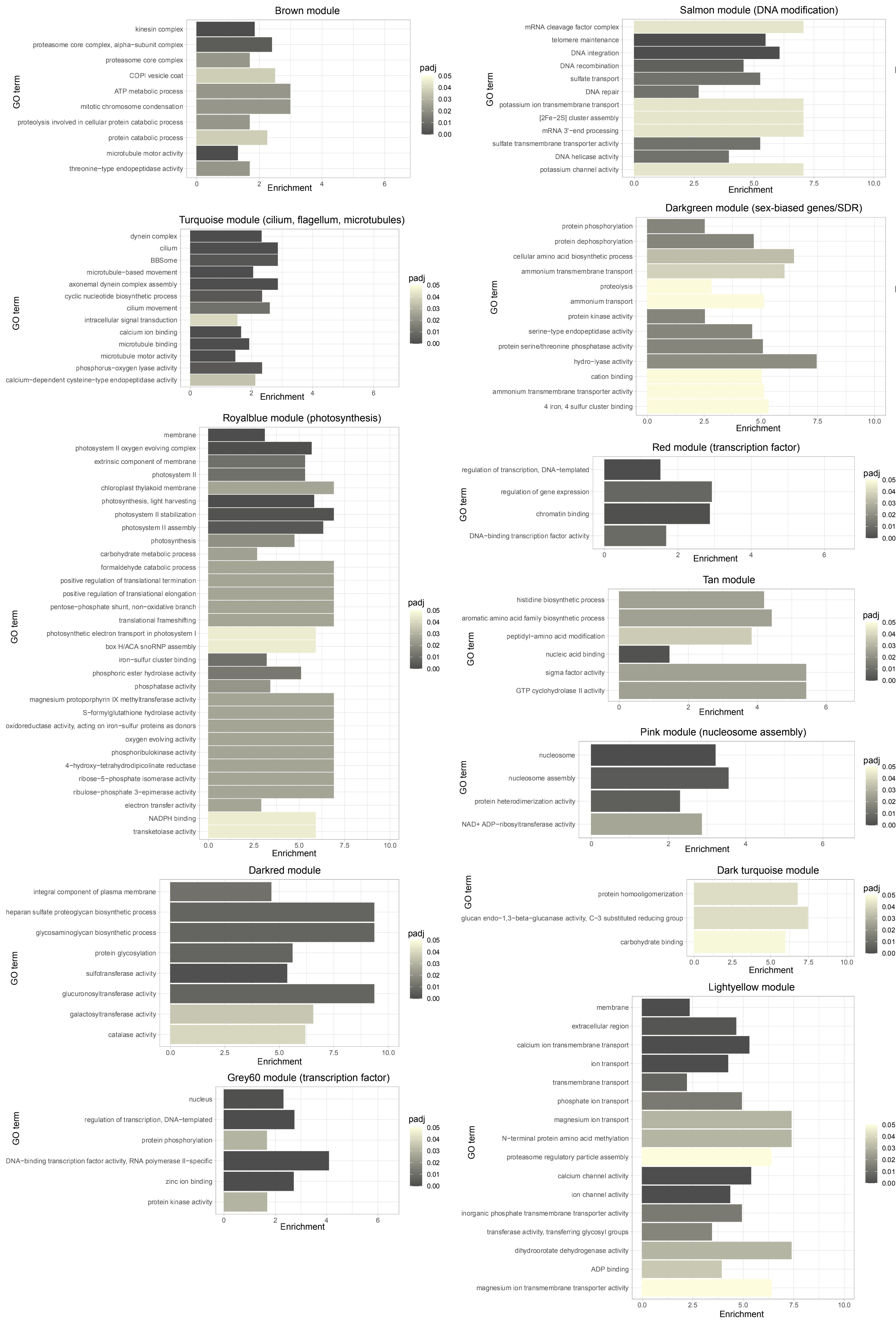

### Fig. S6

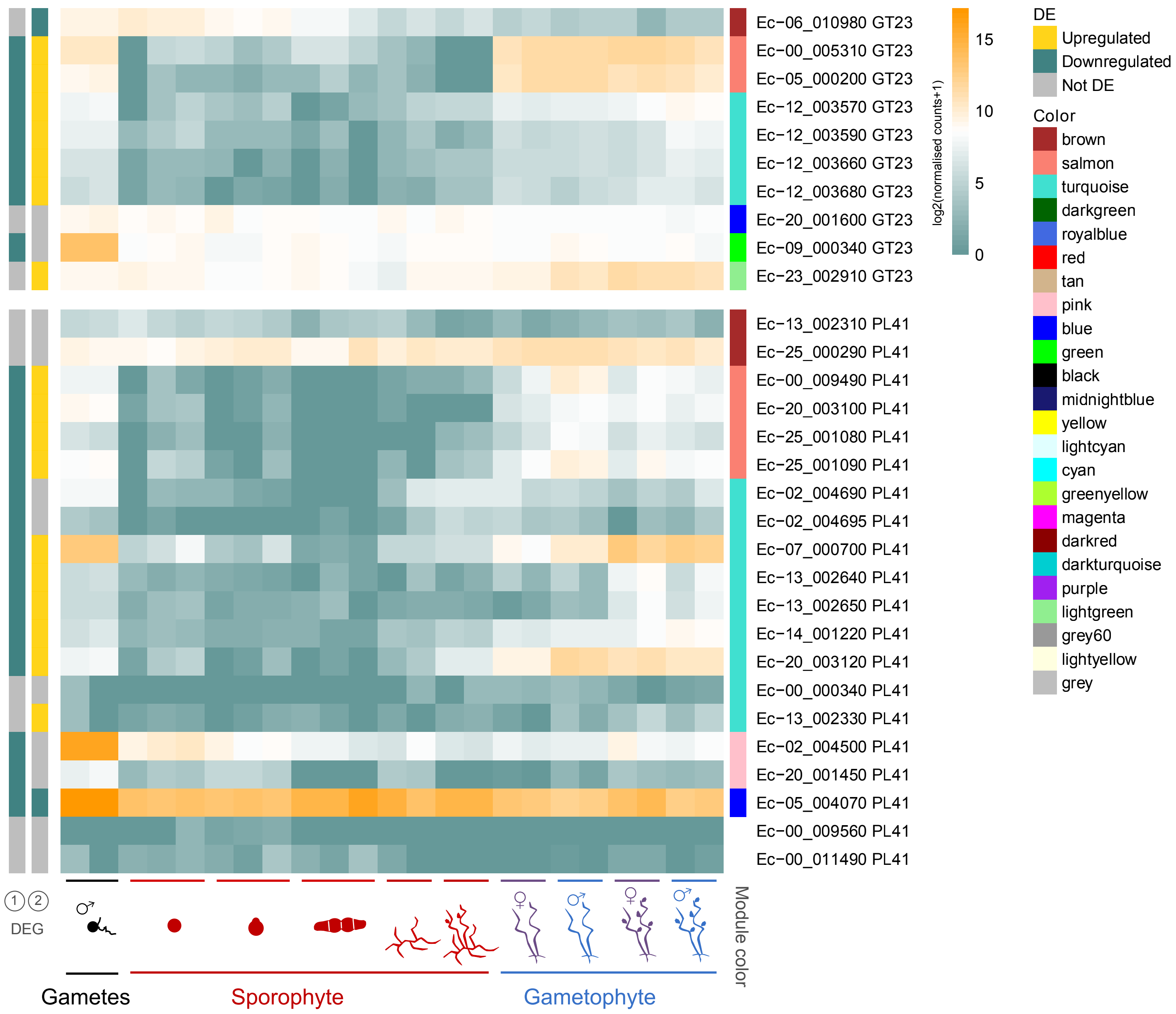

### Fig. S7

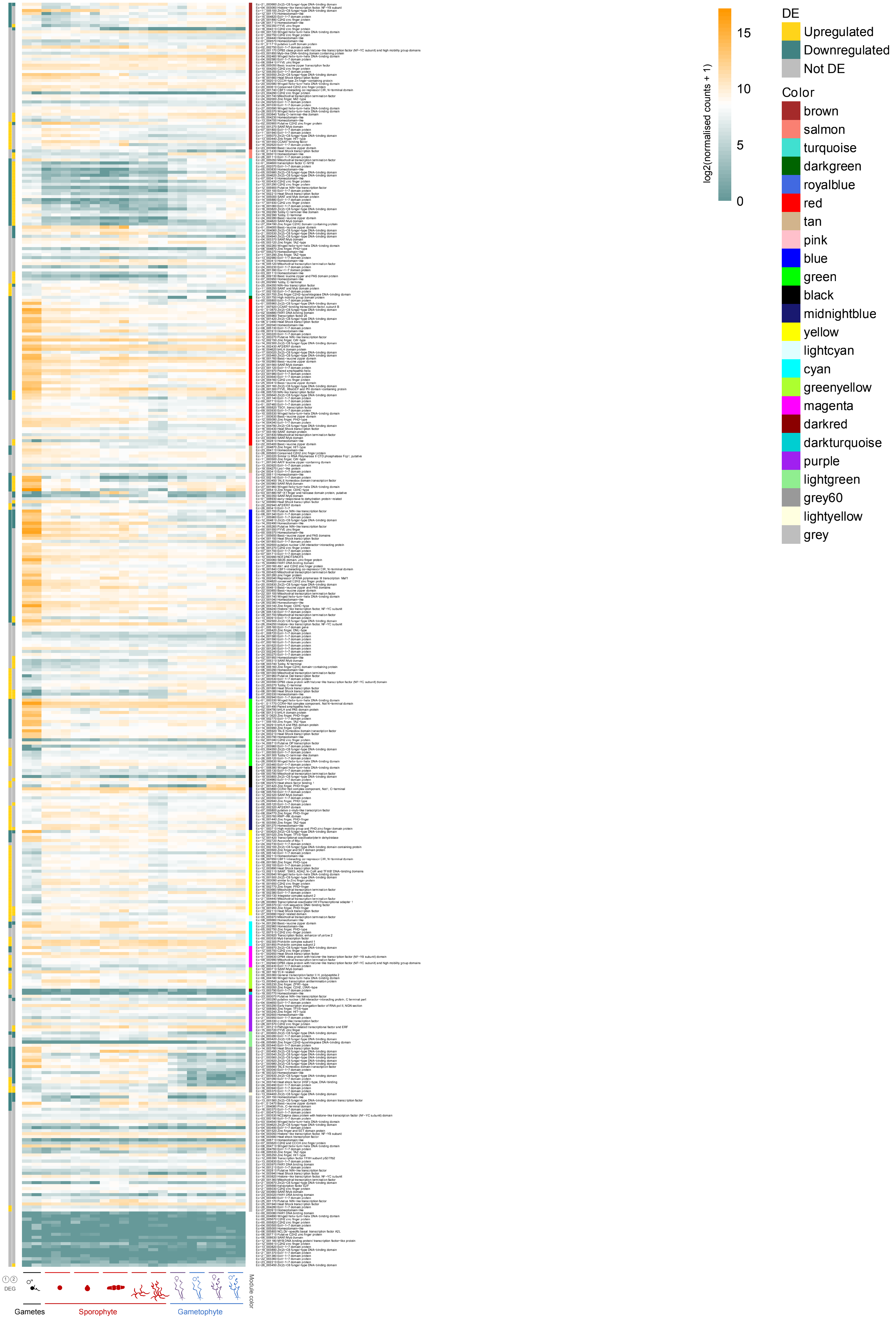

### Fig. S8

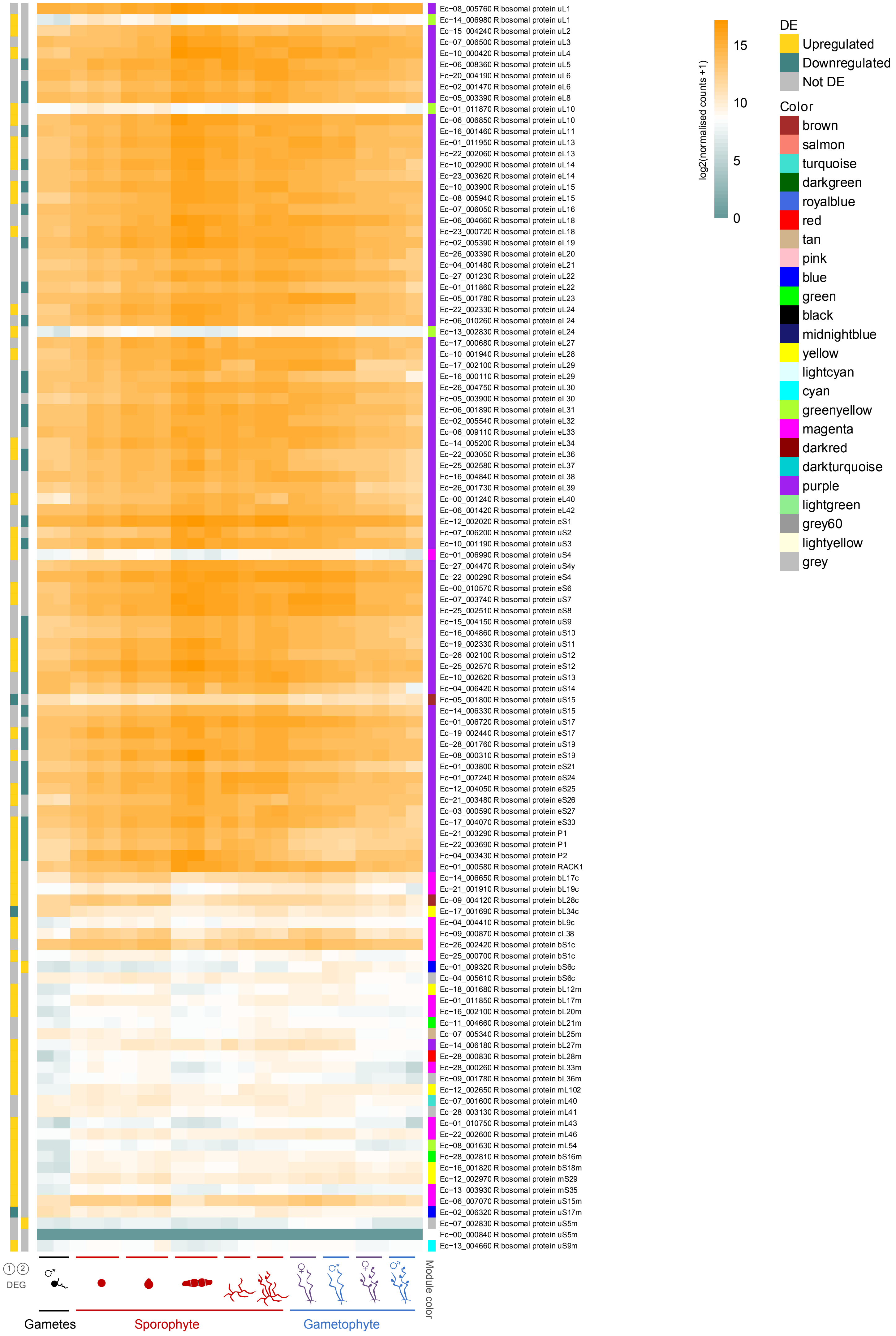

### Fig. S9

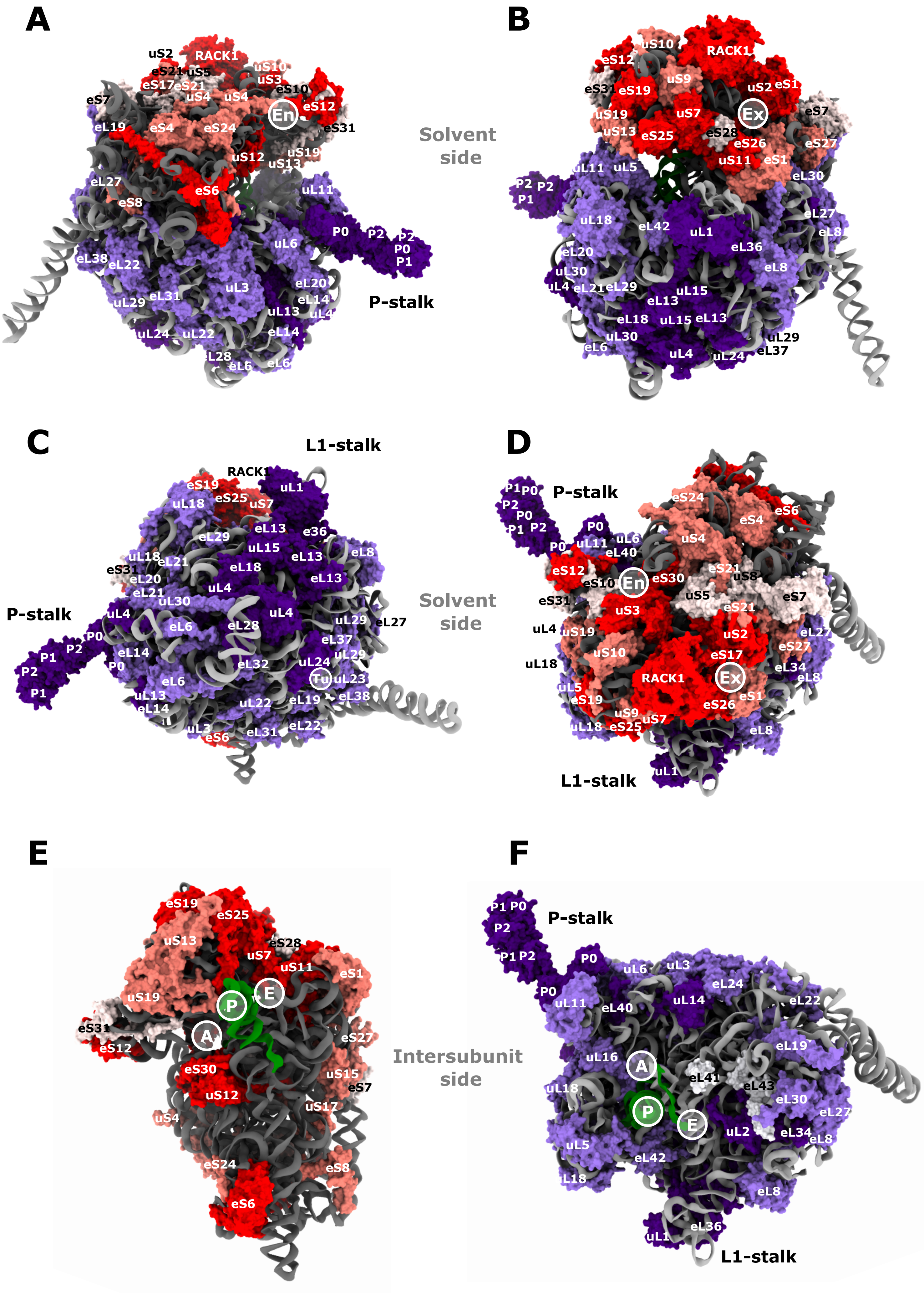

### Fig. S10

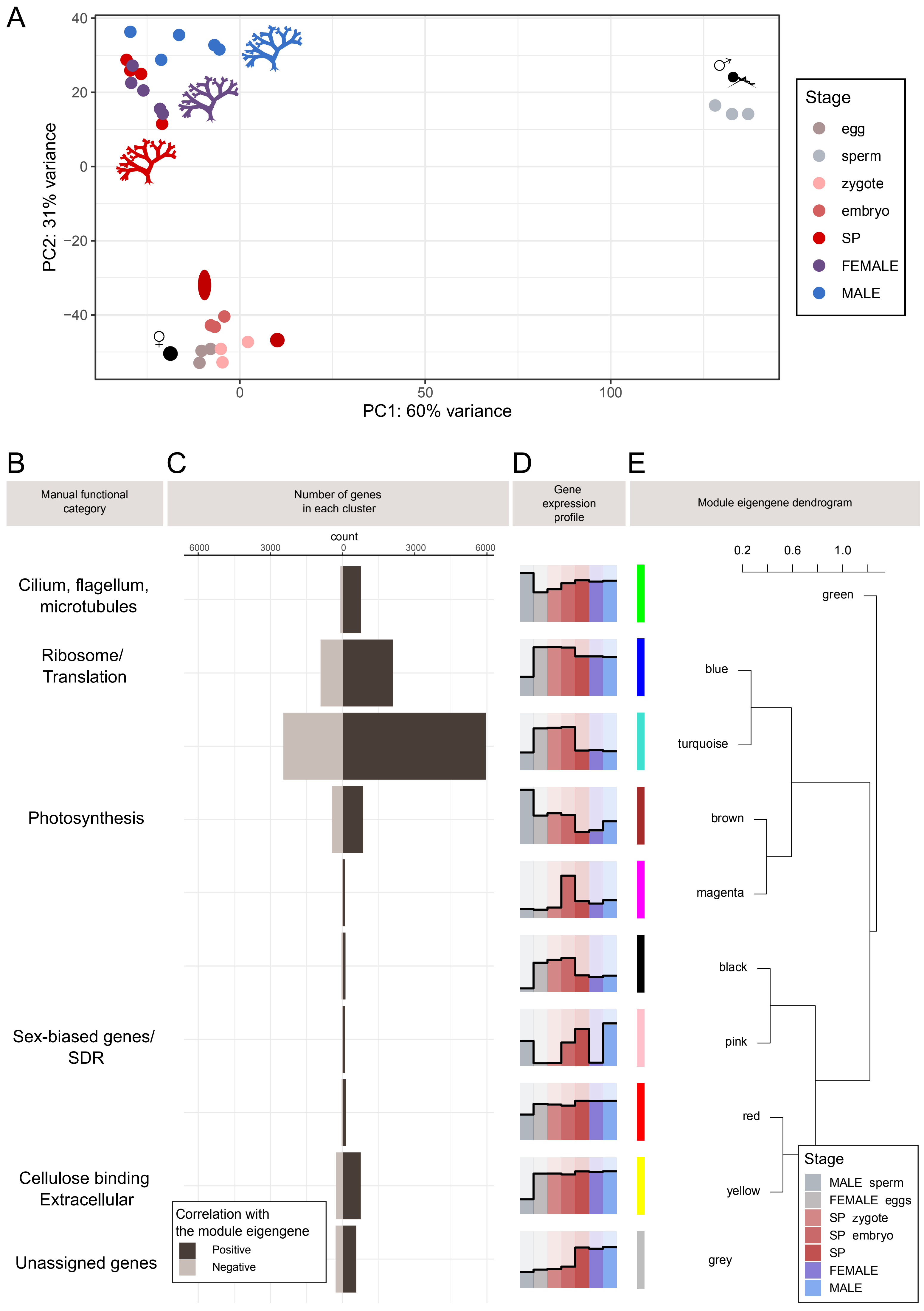

### Fig. S11

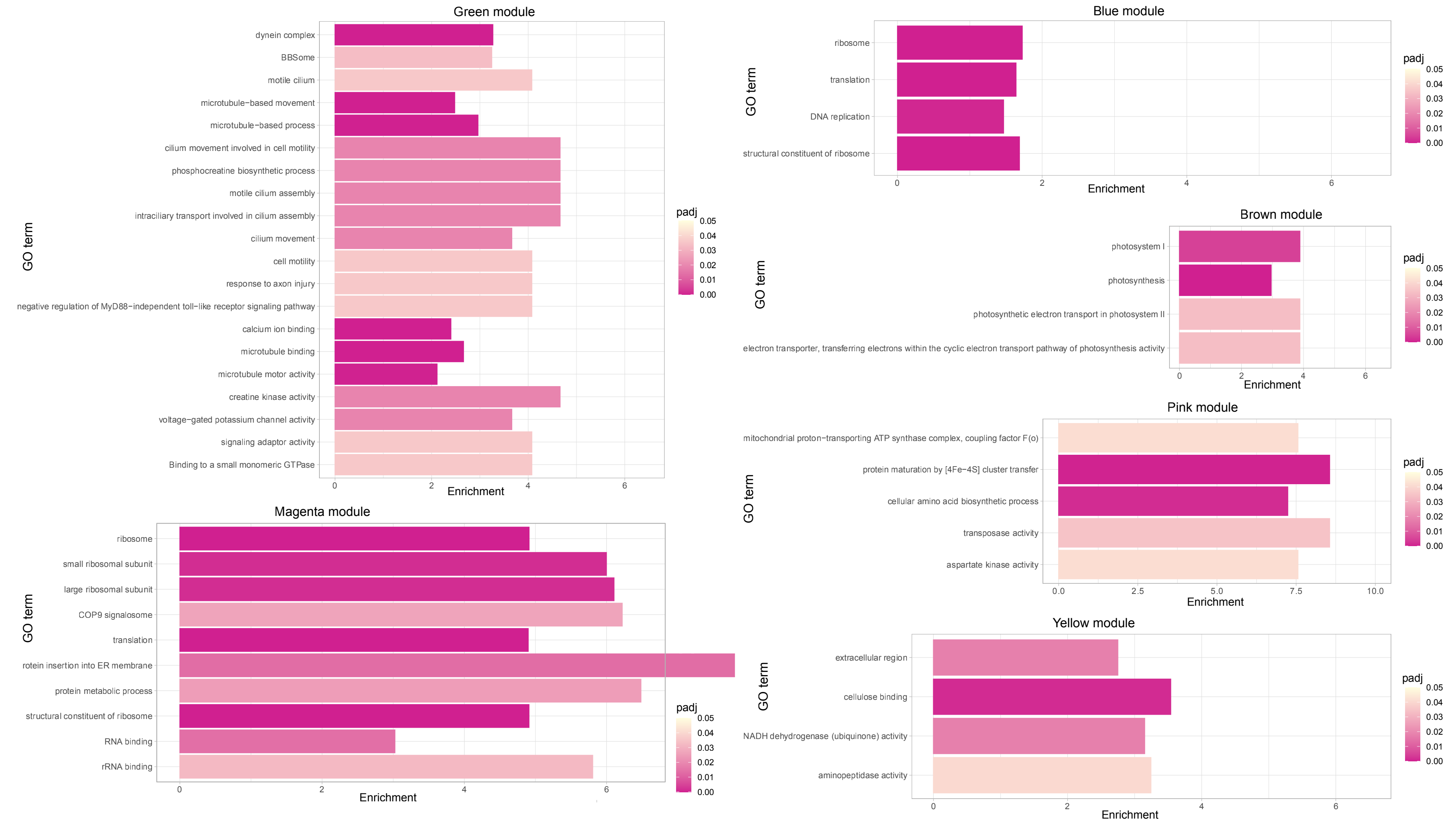

### Fig. S12

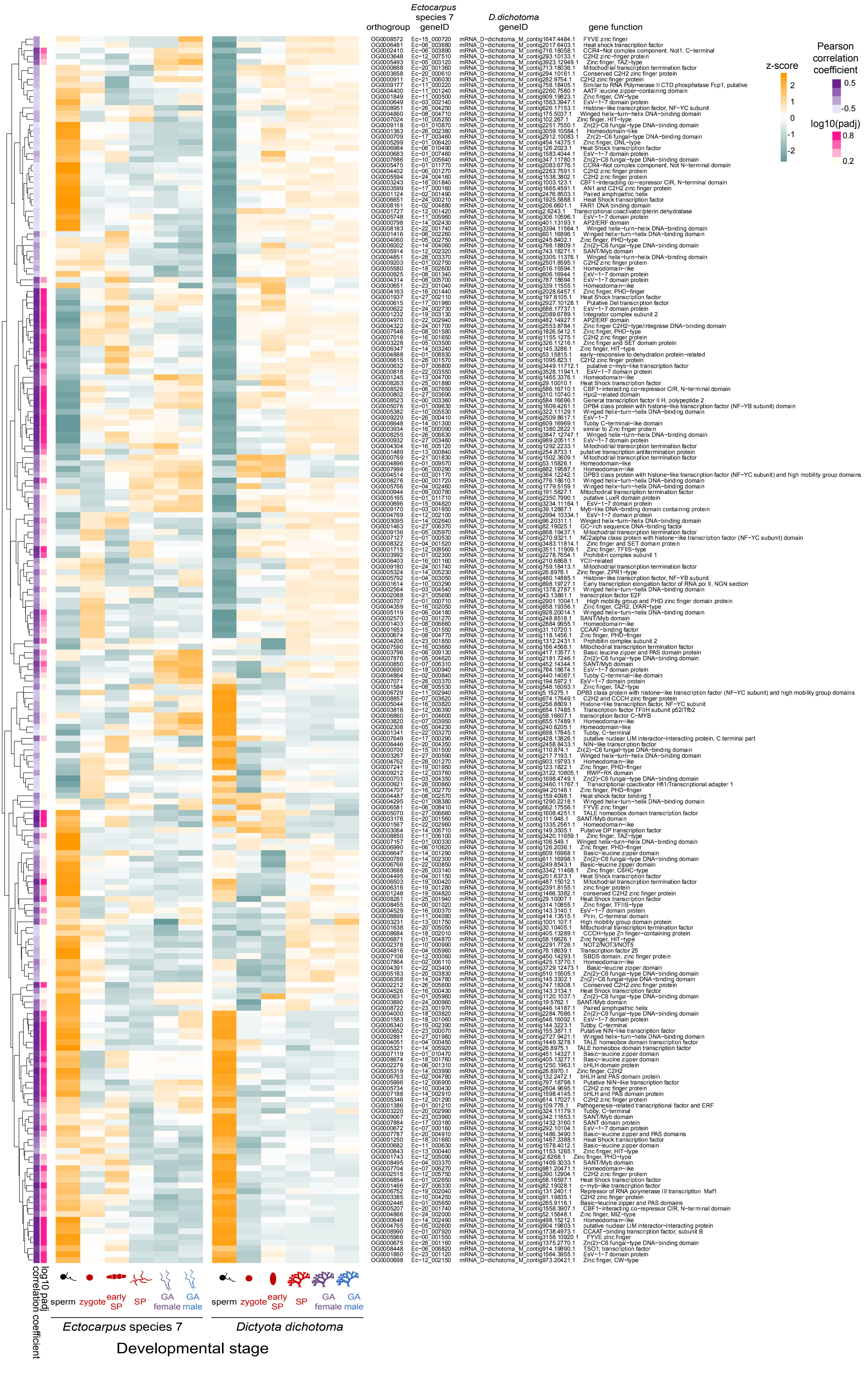
